# Pericytes and Astrocytes Instruct Glioblastoma Invasion, Proliferation, and Therapeutic Response within an Engineered Brain Perivascular Niche Model

**DOI:** 10.1101/2022.04.27.489740

**Authors:** Mai T. Ngo, Jann N. Sarkaria, Brendan A.C. Harley

## Abstract

Glioblastoma (GBM) tumor cells are found in the perivascular niche microenvironment and are believed to associate closely with the brain microvasculature. However, it is largely unknown how the resident cells of the perivascular niche, such as endothelial cells, pericytes, and astrocytes, influence GBM tumor cell behavior and disease progression. We describe a three-dimensional *in vitro* model of the brain perivascular niche developed by encapsulating brain-derived endothelial cells, pericytes, and astrocytes in a gelatin hydrogel. We show that pericytes and astrocytes explicitly contribute to vascular architecture and maturation. We use co-cultures of patient-derived GBM tumor cells with brain microvascular cells to identify a role for pericytes and astrocytes in establishing a perivascular niche environment that modulates GBM cell invasion, proliferation, and therapeutic response. Engineered models provides unique insight regarding the spatial patterning of GBM cell phenotypes in response to a multicellular model of the perivascular niche. Critically, we show that engineered perivascular models provide an important resource to evaluate mechanisms by which inter- cellular interactions modulate GBM tumor cell behavior, drug response, and provide a framework to consider patient-specific disease phenotypes.

## 1. Introduction

Glioblastoma (GBM) is the most common primary malignant brain tumor.^[1]^ Prognosis is generally grim, with a median survival time of approximately one year and a five-year survival rate between 5% and 10%.^[1–2]^ The current standard of care for GBM includes surgical resection followed by treatment with radiation and the alkylating agent temozolomide (TMZ). However, GBM cells are diffusely infiltrative and current radiation and chemotherapy regimens are marginally effective, which leads to invariable recurrence after initial treatment and ultimately patient mortality.^[3]^ Thus, there is a critical need to develop next-generation therapies that can mitigate the invasive capacity and enhance the therapeutic responsiveness of residual cells that remain after surgery.

While cell-intrinsic genomic alterations within GBM play a crucial role in driving tumor cell invasion and therapeutic response,^[4]^ many recent studies also suggest a significant contribution from the tumor microenvironment.^[5]^ In particular, subpopulations of GBM tumor cells have been shown to reside in the perivascular niche (PVN), the local microenvironment adjacent to brain vasculature.^[6]^ The GBM-PVN interactions plays a role in GBM invasion, as tumor cells co-opt and migrate along blood vessels into the surrounding brain tissue.^[7]^ The PVN is also hypothesized to support the activity of cancer stem cells,^[6]^ a subpopulation of tumor cells that are resistant to therapy and are believed to contribute to disease recurrence through self-renewal and differentiation that reproduces the cellular heterogeneity of the original tumor.^[8]^ While it is apparent that the PVN provides a microenvironment that supports disease progression, its explicit role in proliferation and chemoresistance remains underexplored. The precise mechanisms by which the biophysical, biochemical, and cellular components of the PVN direct tumor cell behavior are also poorly developed. Recent studies of the PVN in other types of tumors suggest that signals derived from endothelial and perivascular stromal cells influence phenotypes such as dormancy and chemoresistance.^[9]^ However, these angiocrine signals in GBM remain to be elucidated.

Identifying the mechanisms by which resident perivascular niche cells influence tumor cell behavior is challenging due to the lack of appropriate models. While animal models are the pre-clinical gold standard for therapeutic development, these can be costly and time-intensive to develop for routine experimentation. Whereas therapeutic response in animal models is assessed over a time period spanning weeks to months^[10]^, the ability to accurately assess therapeutic response in a matter of days would enable the ability to test and refine therapeutic regimens for patients in real time. Furthermore, it is challenging to separate the effects of the perivascular niche from other microenvironments also present in the tumor tissue. As a result, there exists an urgent need to develop *in vitro* models that can recapitulate aspects of the perivascular niche environment in order to accelerate mechanistic studies and the evaluation of therapeutic efficacy. A majority of existing *in vitro* models are two-dimensional;^[11]^ while these models have been foundational in identifying potential crosstalk mechanisms between endothelial, perivascular stromal, and tumor cells, they fail to capture the architecture of three-dimensional vasculature, which makes analysis of spatial organization between various cell types within the perivascular niche difficult. Furthermore, while the individual role of endothelial cells, pericytes, and astrocytes have been investigated, synergistic interactions between these cell types likely defines the impact of the perivascular niche on glioblastoma tumor cell phenotype *in vivo*. There is an opportunity for three-dimensional engineered models that capture not only multicellular environment of the perivascular niche, but also the spatial organization of GBM tumor cells within microvascular networks. This approach builds on prior work which showed that endothelial and stromal cells can self-assemble into networks within hydrogels,^[12]^ our adoption of immortalized cell lines and conditioned media assays as a first generation GBM-PVN model,^[13]^ and recent efforts creating a microfluidic model of the brain microvasculature to study metastatic extravasation.^[14]^ Platforms that capture interactions between patient-derived or primary brain tumor cell specimens and an engineered microvasculature network offer the opportunity to assess the impact of angiocrine signals on tumor cell invasion, proliferation, and therapeutic response.

In this study, we report the development and characterization of brain-mimetic microvascular networks within gelatin hydrogels as a three-dimensional model of the brain perivascular niche. We find that brain-derived pericytes and astrocytes play a critical role in shaping the complexity and maturation of microvascular networks. We then describe a co-culture that combines patient-derived glioblastoma tumor cells with perivascular networks comprised of brain microvascular endothelial cells, pericytes, and astrocytes. We leverage the combinatorial nature of this model to identify explicit contributions of pericytes and astrocytes within the perivascular model towards modulating glioblastoma invasion, proliferation, and therapeutic response. Notably, the inclusion of pericytes and astrocytes generate a perivascular environment that globally enhances tumor cell migration and additionally induces spatial patterns of tumor cell behavior, such as increased localization of proliferative GBM cells as well as those harboring stem cell associated markers in close proximity to the perivascular networks. Finally, we use our model to explore the role of pericytes and astrocytes within the perivascular niche in reducing therapeutic response.

## 2. Results

### 2.1. Self-Assembly of Microvascular Networks from Human Brain Vascular Cells in GelMA Hydrogels

We first sought to establish microvascular networks within methacrylamide-functionalized gelatin (GelMA) hydrogels from brain-derived cells. To this end, we cultured human brain microvascular endothelial cells (EC) alone or in combination with human brain pericytes (PC) and astrocytes (AC) to assess the role of pericytes and astrocytes in supporting microvascular network formation in GelMA hydrogels (**Figure 1A**). Cell populations could be individually resolved within multicellular cultures via protein expression: CD31 (endothelial cells), PDGFRβ (pericytes), or GFAP (astrocytes) (**Figure S1A**). We additionally confirmed that human brain microvascular endothelial cells were capable of self-assembling into primitive networks when subjected to a Matrigel tube formation assay (**Figure S1B**). Within GelMA hydrogels, encapsulation of endothelial cells alone (EC), co-cultures of endothelial cells and pericytes (3:1 EC:PC), or tri-cultures containing endothelial cells, pericytes, and astrocytes (3:1:1 EC:PC:AC) resulted in stable microvascular network self-assembly (**Figure 1B**).

**Figure 1.**
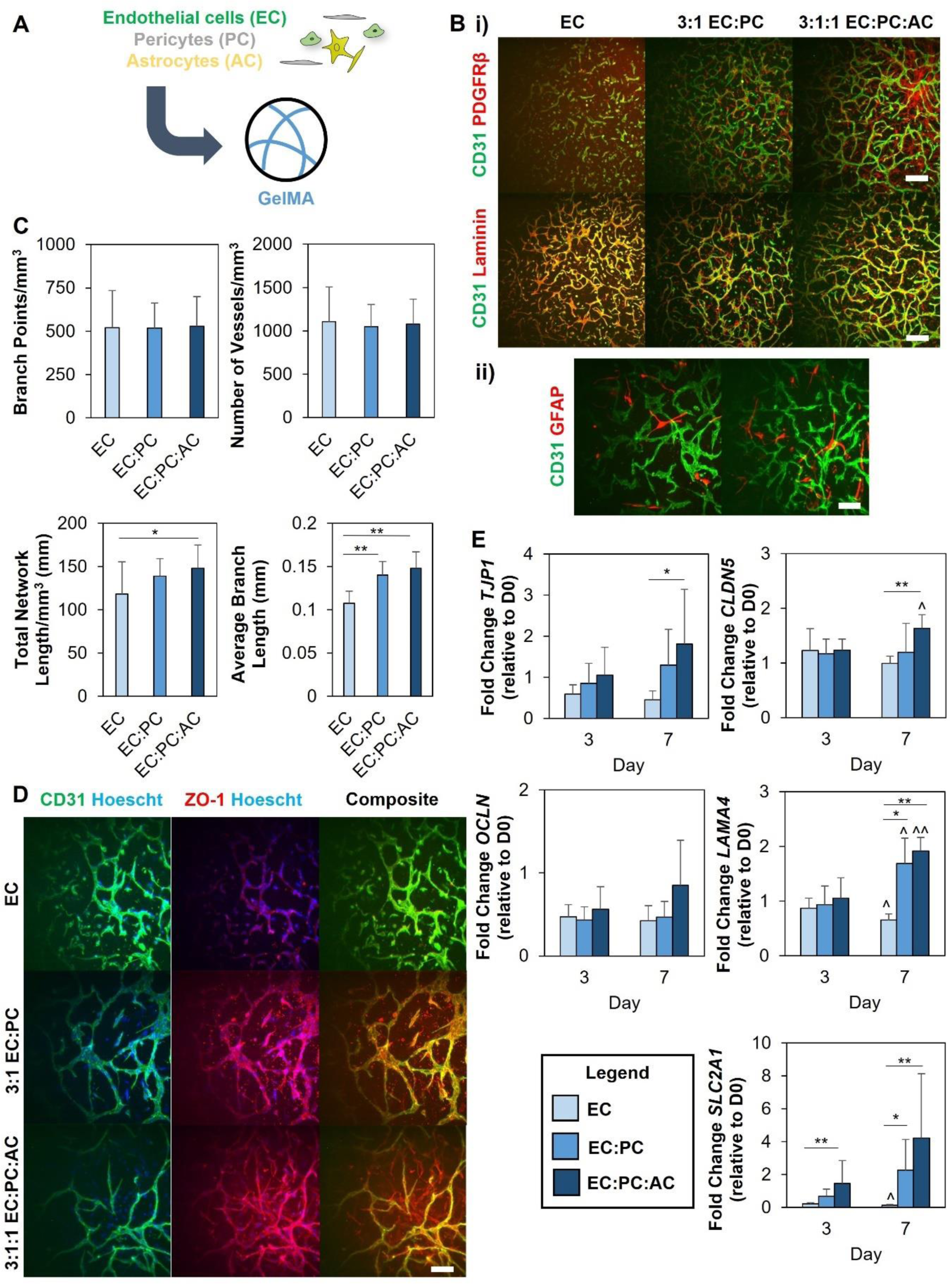
**(A)** Endothelial cells, either alone or combined with pericytes or pericytes and astrocytes, are encapsulated in GelMA hydrogels to self-assemble into microvascular networks. **(B) i)** PDGFRβ expression and laminin deposition in microvascular networks. Scale bar = 200 μm **ii)** GFAP+ astrocytes interacting with microvascular networks in EC:PC:AC cultures. Scale bar = 100 μm **(C)** Quantification of microvascular network architecture. *p<0.05, **p<0.001; N = 15 hydrogels **(D)** Immunofluorescent staining for ZO-1 in microvascular cultures. Scale bar = 100 um **(E)** Gene expression trends for laminin (*LAMA4*), membrane transporter GLUT1 (*SLC2A1*), and tight junction proteins (*TJP1*, *CLDN5*, *OCLN*). *p<0.05, **p<0.01 between groups; ^p<0.05,^^p<0.01 compared to Day 3. N = 6 hydrogels per time point

Notably, addition of pericytes and astrocytes led to increased metrics of microvascular network formation. While there were no significant differences in the number of branch points or vessels between the three culture conditions, there were significant increases in average branch length with the addition of pericytes and astrocytes. Microvascular networks containing endothelial cells, pericytes, and astrocytes (EC:PC:AC) also showed a significant increase in total network length (148 ± 27 mm/mm^3^) compared to endothelial cells (118 ± 37 mm/mm^3^) alone (**Figure 1C**). Immunofluorescent staining also confirmed that PCs assumed a perivascular position to the microvascular network and that astrocytes showed punctate regions of contact or interaction with the microvascular networks (**Figure 1B**).

### 2.2. Presence of Perivascular Stromal Cells Modulates Gene Expression Profile of Microvascular Networks

Having established that we could form brain microvascular networks within GelMA hydrogels, we examined whether these microvascular networks recapitulated other characteristics of *in vivo* brain microvasculature, such as the deposition of basement membrane proteins and the expression of tight junction proteins. Immunofluorescent staining confirmed the perivascular deposition of laminin adjacent to the microvascular network (**Figure 1B**), as well as the localized expression of tight junction proteins ZO-1 and occludins (**Figures 1D** and **S2**). To better compare differences between culture conditions, we used RT-PCR to track and compare gene expression related to tight junctions, basement membrane, and glucose transporters (**Figure 1E**). Amongst the tight junction genes, *TJP1* (which encodes for the protein ZO-1) and *CLDN5* were upregulated by the presence of astrocytes. *LAMA4* expression increased with time in EC:PC and EC:PC:AC networks and was upregulated compared to EC-only cultures. *SLC2A1*, which encodes for the transporter GLUT1, was also upregulated in cultures containing pericytes and astrocytes. Additionally, *LAMA4* and *SLC2A1* expression decreased with time in EC-only cultures that lacked perivascular cell (astrocytes, pericytes) support.

**Figure 2.**
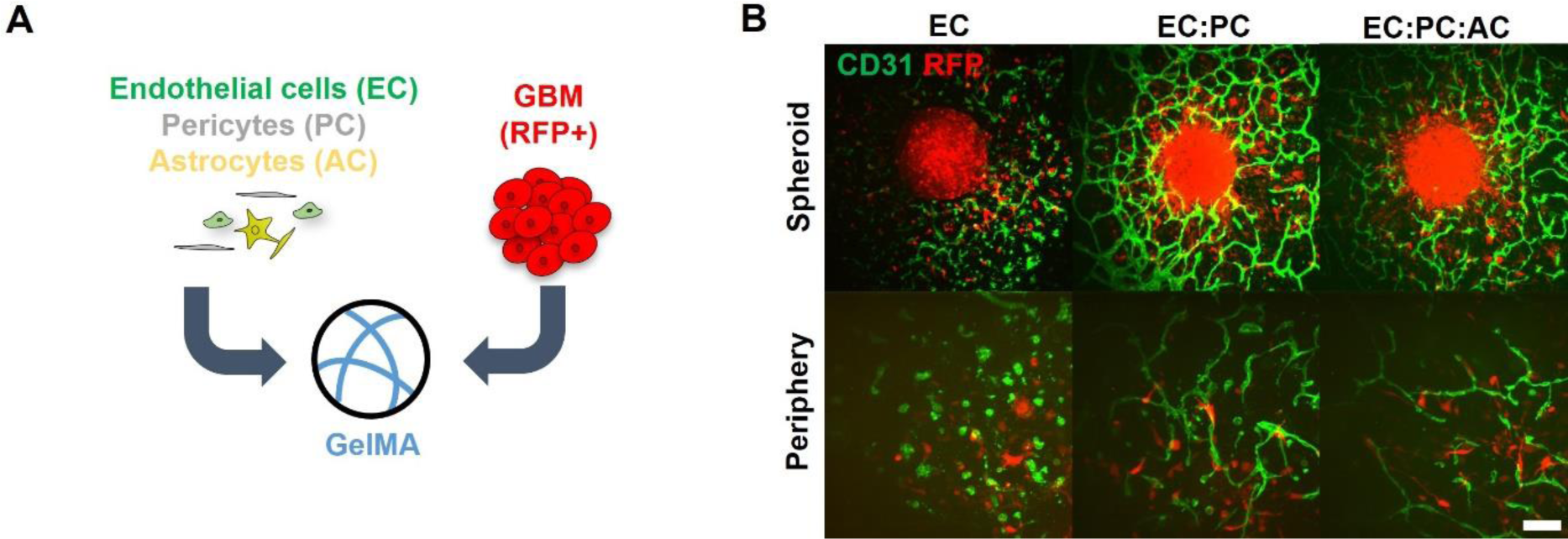
**(A)** Spheroids of RFP+ GBM6 cells are embedded in hydrogels alone or with different combinations of endothelial cells, pericytes, and astrocytes. **(B)** Immunofluorescent staining reveals microvascular network formation surrounding GBM6 spheroids after seven days of culture, and interactions between invading tumor cells and microvasculature in the periphery surrounding the spheroids. Scale bar = 200 μm for spheroid images, 100 μm for periphery images

### 2.3. Combined Presence of Endothelial and Perivascular Stromal Cells Enhance GBM6 Invasion

To then investigate the contributions of brain vascular cells in instructing GBM cell behavior, we generated GBM perivascular niche model consisting of a co-culture of patient-derived xenograft GBM specimens (GBM6 cells, provided by J. Sarkaria, Mayo Clinic) alongside different combinations of the brain vascular cells (EC, EC:PC, or EC:PC:AC). GBM6 cells display a classical subtype, possess *EGFR* amplification and *EGFR*viii mutation, are capable of neurosphere formation, and demonstrate aggressive invasion in orthotopic xenograft models.^[15]^ We confirmed that GBM6 cells cultured using neurosphere conditions expressed neural stem cell markers such as Nestin, Olig2, and Sox2 (**Figure S3A**). These cells retained the ability for multipotent differentiation, as culturing the cells in FBS-containing media led to expression of neural and glial markers βIII tubulin, GFAP, and O1 (**Figure S3B**) as well as a decrease in the expression of neural stem cell markers (**Figure S3C**).

We subsequently developed a spheroid assay to assess the effect of brain microvascular cells on GBM6 invasion (**Figure 2A**). We encapsulated a single spheroid formed from GBM6 cells along with single-cell suspensions containing different combinations of brain vascular cells (ECs only, EC:PC, or EC:PC:AC) in GelMA hydrogels. Hydrogels containing only GBM6 spheroids were used as a baseline control. GBM6 spheroids demonstrated extensive outgrowth into GelMA hydrogels in all conditions over a period of seven days. In hydrogels containing vascular cells, invading GBM6 cells were observed in close proximity to the developing microvascular networks (**Figure 2B**). Beginning at Day 3, spheroids in hydrogels containing the most complex microvascular networks (EC:PC:AC) exhibited significantly larger outgrowth areas compared to hydrogels without vascular cells. By Day 5, spheroids in less complex microvascular networks (EC:PC) also displayed significantly larger outgrowth areas compared to GBM6-only controls (**Figures 3A** and **3B**). No significant invasion advantage was observed for GBM6 cells in the presence of ECs alone relative to GBM6-only conditions. As a complementary approach to assess spheroid invasion, radial profiles of normalized fluorescent intensity were obtained from binarized images of each spheroid, with intensity representing the GBM6 cell density at a radius (r) from the center of the spheroid (**Figure 3C**). Ri metrics, defined as the radius at which there is i% of the maximum intensity, were used to compare the invasive spread of GBM6 cells between the different hydrogel conditions (**Figure 3D**). EC:PC and EC:PC:AC hydrogels displayed the highest r25, r50, and r75 values, which were all significantly higher than the values measured in EC hydrogels. We also observed significant increases in r25 and r50 measurements in EC:PC and EC:PC:AC hydrogels compared to GBM6-only hydrogels. We observed a significant increase in total pixel intensity, indicative of the total number of GBM6 cells, in EC:PC and EC:PC:AC hydrogels compared to GBM6-only and EC hydrogels (**Figure S4**). Normalizing total pixel intensity to outgrowth area, which represents the number of cells per unit area revealed a slight yet significant difference between EC and EC:PC:AC hydrogels (**Figure S4**). Collectively, this data suggests that the incorporation of pericytes and astrocytes into a perivascular niche model significantly enhances the number and invasion distance of tumor cells that invade compared to hydrogels that completely lack vascular cells or co-cultures that only contained endothelial cells alone. The addition of pericytes and astrocytes within a perivascular niche model also increased the apparent cell density within the invasive cohort compared to endothelial cells alone.

**Figure 3.**
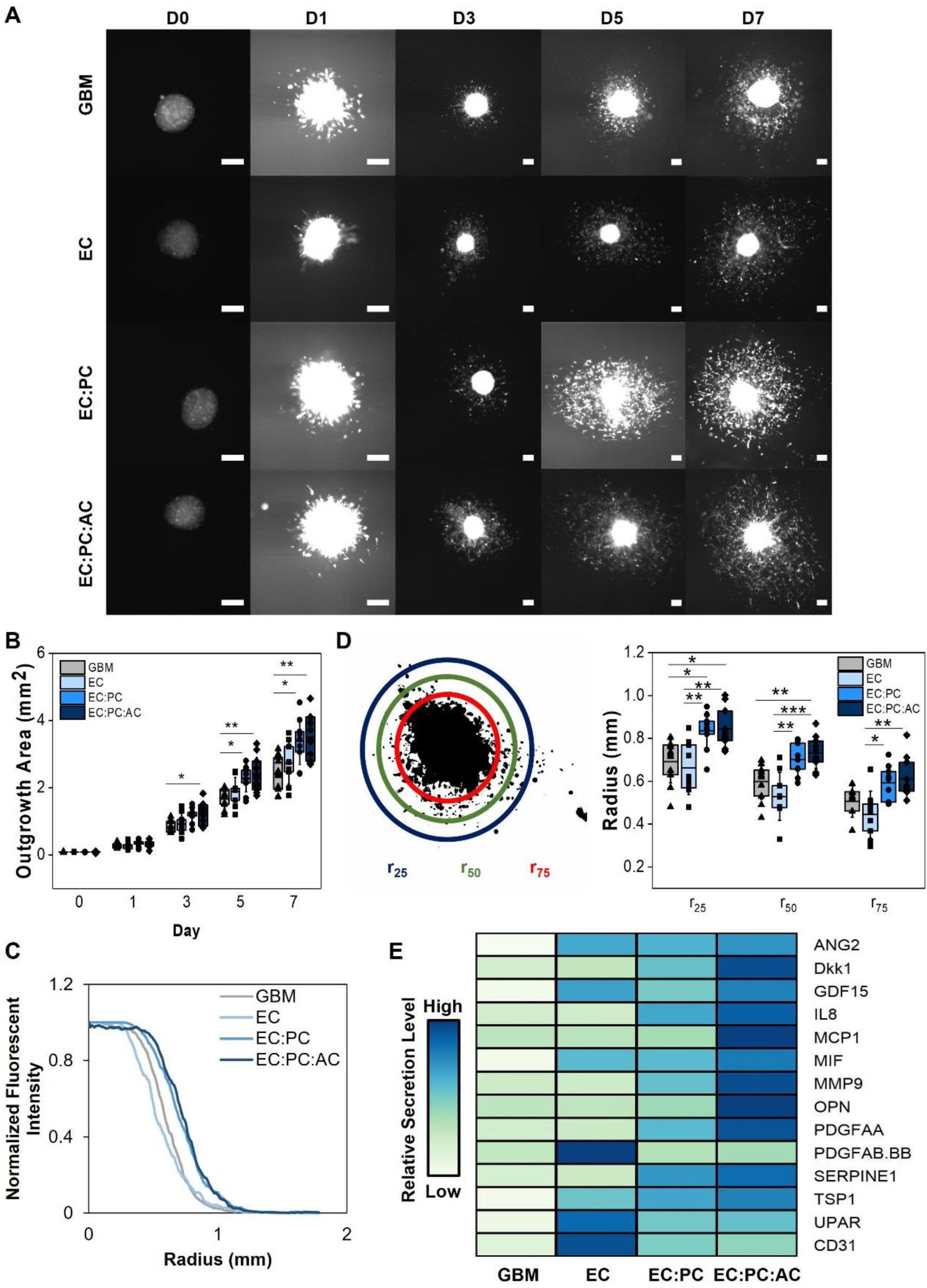
**(A)** Time-lapse images of spheroid outgrowth. Scale bar = 200 μm **(B)** Quantification of spheroid outgrowth area over time. *p<0.05,**p<0.01 between groups **(C)** Radial distribution of fluorescent intensity to quantify tumor cell density as a function of distance away from the spheroid centroid (r = 0) at Day 7. **(D)** ri metrics are defined as the radius at which the fluorescent intensity from RFP-expressing tumor cells is i% of the maximum fluorescent intensity observed in the spheroid core. Data is obtained from Day 7. *p<0.05, **p<0.01, ***p<0.001; N = 9 – 11 hydrogels. **(E)** Heatmap displaying the relative secretion levels of various proteins in spheroid invasion cultures. Secretion levels are normalized for each protein, and therefore should only be compared between groups for a given protein and not between proteins.

### 2.4. Co-Culture of GBM6 Spheroids with Endothelial and Perivascular Stromal Cells Modulates Secretome Profile

We subsequently compared the secretomic profiles for GBM6 spheroids alone versus GBM6 spheroids embedded in EC, EC:PC, or EC:PC:AC perivascular cultures (**Figures 3E** and **S5**) to identify potential paracrine mediators of GBM6 invasion. Compared to GBM6 spheroids alone, the inclusion of ECs resulted in increased secretion of ANG2, GDF-15, PDGFAB/BB, UPAR, and CD31. The inclusion of additional perivascular stromal cells (astrocytes and pericytes) led to significant increases in the secretion of Dkk1, IL-8, MCP1, MIF, MMP9, OPN, PDGFAA, TSP1, and SERPIN E1. We subsequently cross-referenced these proteins against the Cancer Genome Atlas glioblastoma dataset using GlioVis to identify corresponding genes upregulated in human glioblastoma specimens compared to normal brain tissue,^[4a, 16]^ finding a critical group of factors (ANG2, GDF-15, MMP9, OPN, SERPIN E1, UPAR, CD31) upregulated in glioblastoma and generated by engineered perivascular cultures.

### 2.5. Combined Presence of Endothelial and Perivascular Stromal Cells Modulates SOX2 Expression and Proliferation of GBM6 Cells

Having established a role for pericytes and astrocytes in GBM invasion within an engineered perivascular niche, we hypothesized that perivascular stromal cells may impact other facets of GBM cell behavior, such as proliferative capacity and the maintenance of stem cell markers. Thus, we examined expression of SOX2, a pluripotency marker used to identify tumor cells with stem-like behavior,^[17]^ as well as KI67 expression and EdU incorporation, which collectively provide information regarding the cycling or proliferative state of the tumor cells. To more clearly study the role of PVN signaling on GBM cells, we encapsulated single cell distributions of GBM6 specimens (2.5 × 10^5^/mL) within microvascular networks (ECs, EC:PC, or EC:PC:AC networks) rather than cell spheroids. After seven days, not only were robust microvascular networks observed in EC:PC and EC:PC:AC cultures, but the inclusion of pericytes and astrocytes increased the association between tumor cells and the microvascular network (**Figure 4A**). Specifically, the presence of pericytes and astrocytes significantly increased the number of tumor cells within 50 μm of the resulting microvascular network – where 50 μm has been used in prior literature to define the perivascular niche environment.^[9a, 18]^ While a majority (> 70%) of tumor cells expressed SOX2 regardless of the complexity of the perivascular network, we observed reduced SOX2 expression in GBM cells in perivascular cultures that included astrocytes versus those derived only with ECs (**Figures 4B** and **4C**). Interestingly, SOX2+ tumor cells resided closer to the microvascular networks compared to SOX2- tumor cells regardless of the complexity of the perivascular culture (**Figure 4D**). Further, inclusion of pericytes and astrocytes in the perivascular niche model significantly decreased the fraction of KI67-expressing tumor cells compared to GBM6-only cultures, suggesting an increased presence of quiescent cells (**Figures 5A** and **5B**). In hydrogels containing perivascular stromal cells, KI67+ tumor cells also resided closer to the microvascular networks compared to KI67- tumor cells (**Figure 5C**). We subsequently evaluated the fraction of tumor cells that incorporated EdU in a 24-hour pulse starting after 7 days of culture, meaning that EdU-positive went through S-phase of the cell cycle during that 24 hour pulse (**Figure S6**).^[19]^ Notably, we observed a significant increase in the EdU+/KI67+ fraction ratio in the presence of perivascular stromal cells compared to tumor cells alone (**Figure 5D, E**). This data suggests that while GBM cells showed increased overall quiescent (Ki67-) fraction in the presence of perivascular stromal cells, the cells that were KI67+ were much more likely to be actively dividing within a 24-hour period (EdU+). While there was a trend towards EdU+ tumor cells residing more closely to the microvascular networks compared to EdU- tumor cells, the effect was only significant in cultures containing only endothelial cells; EdU+ GBM6 cells were also significantly less likely to be more than 50μm from the microvascular network in models containing perivascular stromal cells (EC:PC:AC; **Figure 5F**). Overall, this data suggests that signals from a perivascular niche environment comprised of both endothelial cells and perivascular stromal cells strongly influences patterns Data points represent the fraction of tumor cells that are a specified distance from the microvascular network in a single hydrogel. *p<0.05, **p<0.01, ***p<0.001; N = 5 - 6 hydrogels of GBM cell quiescence, proliferation, and retention of SOX2 expression in both a context- and location-dependent fashion.

**Figure 4.**
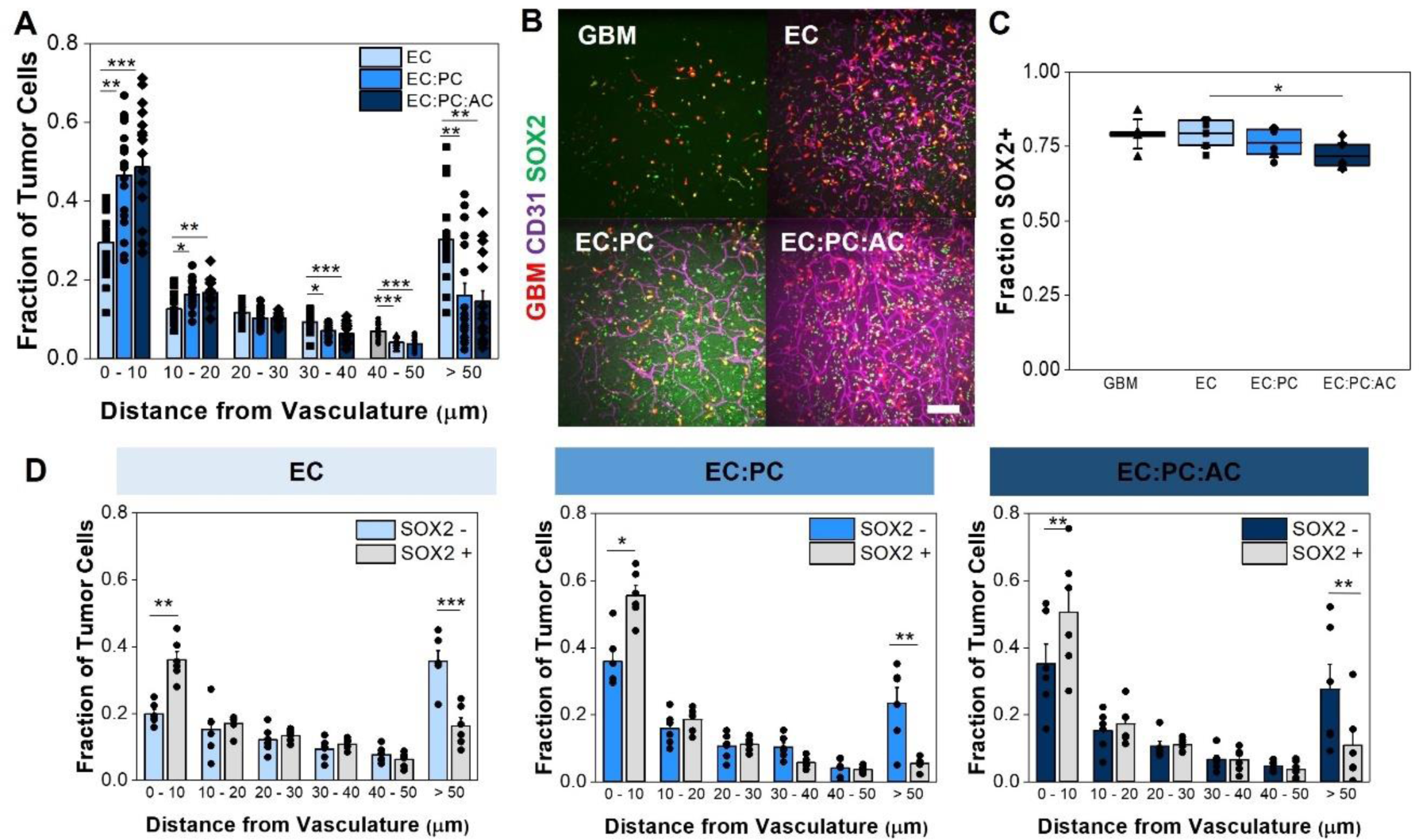
**(A)** Proximity of tumor cells to the microvascular network. Data points represent the fraction of tumor cells that are a specified distance from the microvascular network in a single hydrogel. *p<0.05, **p<0.01, ***p<0.001; N = 17-18 hydrogels **(B)** Representative images of tumor-vascular co-cultures with SOX2 staining. Scale bar = 200 μm **(C)** Quantification of the fraction of RFP-expressing tumor cells that express SOX2. *p<0.05, **p<0.01; N = 6 hydrogels **(D)** Comparing the proximity of SOX2- vs. SOX2+ GBM6 cells to the microvascular network. Data points represent the average distance between a tumor cell and the microvascular network in a single hydrogel. *p<0.05, **p<0.01, ***p<0.001; N = 6 hydrogels

**Figure 5.**
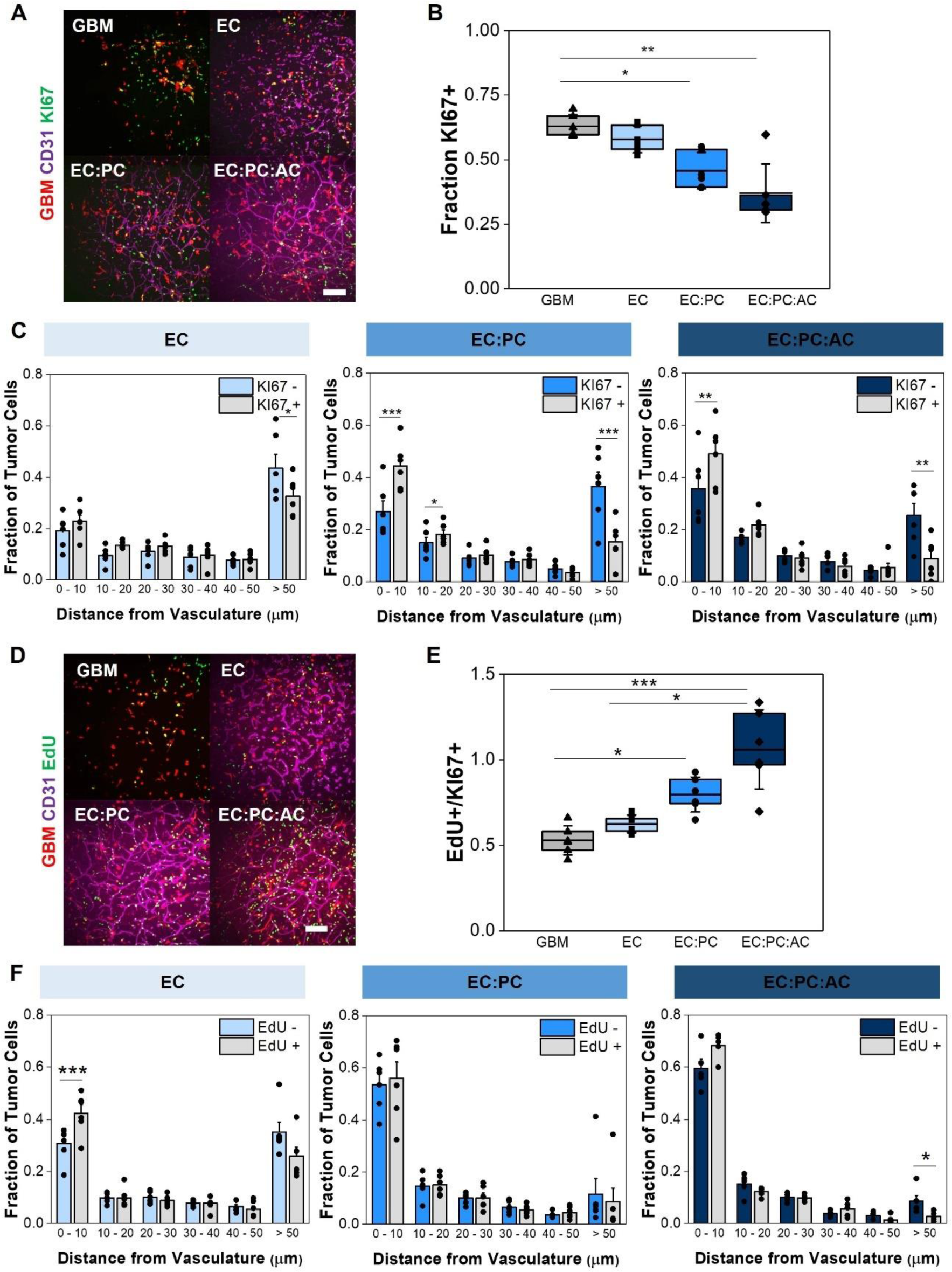
**(A)** Representative images of tumor-vascular co-cultures with KI67 staining. Scale bar = 200 μm **(B)** Quantification of the fraction of RFP-expressing tumor cells that express KI67. *p<0.05, **p<0.01; N = 6 hydrogels **(C)** Comparing the proximity of KI67- vs. KI67+ GBM6 cells to the microvascular network. Data points represent the fraction of tumor cells that are a specified distance from the microvascular network in a single hydrogel. *p<0.05, **p<0.01, ***p<0.001; N = 6 hydrogels **(D)** Representative images of tumor-vascular co- cultures with EdU staining. Scale bar = 200 μm **(E)** Quantification of the fraction of KI67+ tumor cells that incorporate EdU during a 24-hour pulse. *p<0.05, **p<0.01; N = 6 hydrogels **(F)** Comparing the proximity of EdU- vs. EdU+ GBM6 cells to the microvascular network.

### 2.6. Perivascular Stromal Cells Reduce GBM12 Apoptosis in Perivascular Niche Environment and Modulate Proliferation After Temozolomide Treatment

Finally, we wanted to explore contributions of the perivascular niche environment in modulating GBM cell response to temozolomide (TMZ), which is the standard-of-care chemotherapeutic agent for GBM. Because GBM6 cells contain an unmethylated *MGMT* promoter that inherently limits their response to TMZ in biomaterial cultures,^[20]^ we used cells from a separate patient-derived xenograft, GBM12, for our TMZ studies. Similar to GBM6, GBM12 cells have a classical subtype, possess *EGFR* amplification, and invade within orthotopic xenograft models.^[15]^ However, the *MGMT* promoter of GBM12 is methylated,^[21]^ and GBM12 cells are much more sensitive to TMZ. For these experiments, we cultured single-cell suspensions of GBM12 cells in hydrogels on their own or in the presence of our series of perivascular models of increasing complexity (EC, EC:PC, EC:PC:AC). We cultured the hydrogels for seven days before exposing them to 600 µM temozolomide or DMSO as a vehicle control for 48 hours (**Figure 6A**). We chose 600 μM because it was identified as the concentration at which growth was reduced by 50% in our hydrogels (**Figure S7**).^[22]^ Using immunofluorescent staining, we observed increased cPARP^+^ (cleaved PARP) apoptotic cells across all experimental groups compared to DMSO control (**Figures 6B** and **S8**).

**Figure 6.**
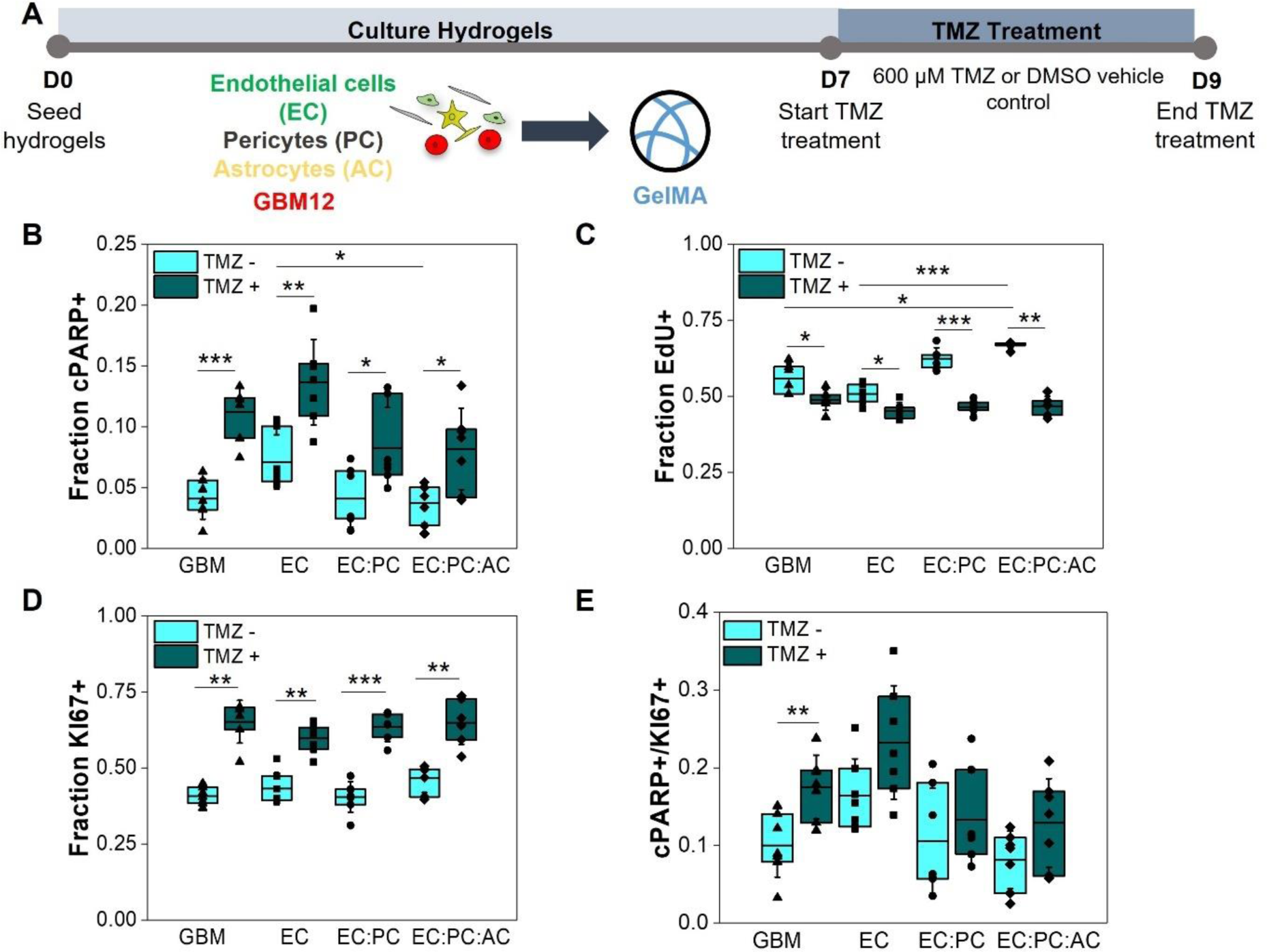
**(A)** Experimental set-up to study tumor cell response to temozolomide or a DMSO control in tumor-only or tumor-vascular cultures. **(B)** Quantification of the fraction of RFP- expressing GBM12 tumor cells that express cPARP with and without temozolomide treatment. **(C)** Quantification of the fraction of RFP-expressing GBM12 tumor cells that incorporate EdU during a 24-hour pulse with and without temozolomide treatment. **(D)** Quantification of the fraction of RFP-expressing GBM12 tumor cells that express KI67 with and without temozolomide treatment. **(E)** The apoptosis-to-proliferation ratio cPARP+/KI67+ for GBM12 tumors with and without temozolomide treatment. *p<0.05, **p<0.01, ***p<0.001. N = 6-7 hydrogels for KI67 experiments, 7 hydrogels for cPARP experiments and cPARP+/KI67+ ratios, and 6 hydrogels for EdU experiments

Interestingly, in DMSO control groups, the number of apoptotic tumor cells was reduced in hydrogels containing pericytes and astrocytes compared to endothelial cells alone. EdU incorporation also decreased in all experimental groups treated with temozolomide (**Figures 6C** and **S9**), while KI67 expression increased with temozolomide treatment across all experiment groups (**Figures 6D** and **S10**). We subsequently examined cPARP+/KI67+ (apoptosis-to-proliferation) ratios as a measure of therapeutic sensitivity.^[23]^ While the cPARP+/KI67+ ratio increased significantly for GBM12 only cultures in response to temozolomide treatment, no significant changes were observed in hydrogels that incorporated perivascular stromal cells (**Figure 6E**), suggesting the perivascular microenvironment can contribute to an environment that protects GBM12 cells from the effects of temozolomide.

## 3. Discussion

Glioblastoma cells interact significantly with perivascular niche environments within the brain,^[6]^ where angiocrine signals have the potential to significantly modulate tumor cell behavior. Several studies have revealed that shifts in extracellular hypoxia^[24]^ and endothelial-mediated signaling influence tumor cell invasion and therapeutic resistance,^[25]^ but the contributions of perivascular stromal cells such as pericytes and astrocytes have not been extensively explored. Pericytes have primarily been studied in the context of tumor angiogenesis, in which they are recruited to developing vasculature and promote vessel stabilization and maturation.^[26]^ Astrocytes contribute to emergence of the blood-brain barrier phenotype;^[27]^ while *in vitro* co-culture studies with glioblastoma tumor cells suggest that signaling from astrocytes increases tumor cell migration, proliferation, and therapeutic resilience, these studies were not performed in the context of the perivascular environment.^[11b, 11c, 28]^ Thus, the effects of synergistic signaling from endothelial cells, pericytes, and astrocytes on glioblastoma behavior in fully three-dimensional models of the brain microenvironment have yet to be defined. Understanding how perivascular stromal cells shape the development and maturation of the perivascular niche, and whether their influence is needed for the perivascular niche to instruct tumor cell behavior, will reveal important insights into the mechanisms that govern microenvironmental regulation of disease progression.

We first established a three-dimensional model of brain microvasculature to recapitulate aspects of the organization and architecture of the perivascular niche. Three-dimensional blood-brain barrier models are an area of recent innovation,^[29]^ and recent work by *Campisi et al.* showed a human brain microvascular culture formed via self-assembly of iPS-derived endothelial cells, pericytes and astrocytes in fibrin gels within a microfluidic device.^[14]^ Inspired by this work, we demonstrate that human brain microvascular networks can be generated in macroscale GelMA hydrogels. The macroscale dimensions of our platform are advantageous for collecting larger sample volumes for downstream analyses such as transcriptomics, proteomics, and secretomics. While many three-dimensional BBB models are generated in collagen or fibrin hydrogels,^[14, 30]^ our use of GelMA enables improved user control over biophysical and biochemical properties via manipulation of UV photopolymerization parameters and chemical modification of the methacrylate backbone.^[31]^ We observed self-assembly of brain microvascular endothelial cells into networks; furthermore, addition of pericytes and astrocytes results in appropriate perivascular positioning of pericytes along the vascular network, while astrocytes extend processes to contact the endothelial networks at distinct locations. Our results reveal important contributions of pericytes and astrocytes towards shaping microvascular architecture and increasing gene expression related to tight junctions, basement membrane proteins, and membrane transporters, which are critical components of a maturing microvascular network. Our gene expression trends align with those reported by *Campisi et al*.,^[14]^ while we report an increase in average branch length with the addition of perivascular stromal cells that contrasts with their observations of a decreasing trend. Overall, we demonstrate that brain-derived pericytes and astrocytes actively contribute to microvascular development and phenotype.

While the absence of perfusion is a current limitation of our model and is an area of ongoing work, GBM cells interact with and spread along vessels rather than intravasating within the vasculature. Thus, while perfusable vessels are important for studying drug distribution and vascular remodeling in response to therapy^[32]^, our model is able to capture the interactions between GBM cells with the external vessel surface and is well-positioned to investigate processes of brain microvascular morphogenesis, dysregulation during pathophysiological conditions, and angiocrine-derived crosstalk with brain parenchymal cells during homeostasis, regeneration, and disease.

Diffuse invasion of tumor cells is a significant clinical challenge in treating glioblastoma,^[33]^ with tumor cells observed to migrate towards and along vasculature as a means of dissemination.^[7a, 34]^ Identifying angiocrine mediators of tumor cell invasion provides potential therapeutic strategies for mitigating invasion. By adapting a spheroid-based invasion assay commonly used in our lab,^[35]^ we investigated the effects of synergistic crosstalk between endothelial cells, pericytes, and astrocytes on tumor cell invasion. In contrast to other reports,^[25c, 36]^ the presence of endothelial cells alone was not sufficient to increase invasion.

These differences may be attributed to varying endothelial cell sources, choice of biomaterials, and culture conditions, which collectively may influence the initial angiocrine state of the endothelial cells and their ability to generate a pro-invasive niche. Regardless, we find that the addition of pericytes and astrocytes alongside endothelial cells forms a more complex perivascular microenvironment that strongly induces a pro-invasive response. While the role of pericytes beyond contributions to angiogenesis have not been explored,^[37]^ results by *Herrera-Perez et al.* and *Rath et al.* support a pro-invasive role for astrocytes.^[11b, 38]^ Inclusion of pericytes and astrocytes provided a signaling environment that decreased the average distance between tumor cells and microvasculature; this finding recapitulates the close association or co-option of vessel by glioblastoma tumor cells *in vivo*.^[7b]^ We compared the secretome generated by different perivascular networks to identify potential paracrine mediators for an enhanced invasive response. Several proteins were consistent with those previously identified in our first-generation perivascular niche platform comprised of human umbilical vein endothelial cells and normal human lung fibroblasts, suggesting a conserved role in capillary morphogenesis regardless of vascular cell source.^[13b]^ Amongst proteins with increased secretion due to the incorporation of perivascular stromal cells, ANG2, SERPINE1, AND TSP1 have been shown to be expressed in vascularized regions of glioblastoma tissue, with ANG2 additionally acting as a marker for vessels co-opted by tumor cells.^[39]^

Furthermore, GDF-15, IL-8, MCP1, MIF, OPN, PDGFAA, and TSP1 have been shown to increase glioblastoma cell invasion via exogenous stimulation or a chemotactic effect.^[40]^ MMP9 was additionally upregulated with the presence of astrocytes, and has been shown to be expressed in vascular regions of glioblastoma tissue and promotes tumor cell invasion.^[41]^ The role of MMP9 in matrix remodelling suggests future studies to probe for biochemical and biophysical changes to the hydrogel microenvironment that may also facilitate tumor cell invasion. Based on our secretomic analysis and by using microfluidic invasion models we have previously described,^[13b, 42]^ ongoing work is studying the individual and combined role of these specific soluble proteins on pro-invasive response of the GBM cells within our model, as well as identifying the cell types that are secreting these proteins. These findings suggest that a complex perivascular niche model assembled from brain-derived endothelial cells along with pericytes and astrocytes not only replicates structural elements of the brain perivascular microenvironment, but also expresses soluble factors consistent with the GBM tumor microenvironment known to enhance GBM cell invasion.

The role of the perivascular niche on GBM cell proliferation is unclear, with some studies demonstrating a proliferative effect from endothelial crosstalk while other studies suggest that crosstalk has no effect on proliferation.^[25c, 43]^ While these prior studies present a population-level analysis of tumor cell proliferation, tumor cells within the perivascular niche reside at varying distances in relation to the microvasculature; this heterogeneity in spatial proximity has been shown to impact phenotypes such as dormancy and migration.^[9a, 18]^ A particular advantage of our platform is the ability to interrogate the effects of spatial proximity between tumor cells and microvasculature on GBM cell behavior by using immunofluorescent staining and image analysis pipelines. Strikingly, we discovered that while the percentage of KI67+ tumor cells decreased in cultures containing perivascular stromal cells, the remaining KI67+ fraction localized signficantly closer to microvascular networks that included perivascular stromal cells. These results suggest that paracrine versus juxtacrine signals and potential matrix remodeling events in the perivascular space impact tumor cell proliferation differentially. Notably, the observed dichotomy between enhanced invasion and decreased KI67 expression broadly correlates to the “go-or-grow” hypothesis that suggests that migration and proliferation are mutually exclusive events in glioblastoma.^[7a, 44]^ These findings are also consistent with our prior RNA-sequencing results that found that tumor-vascular cell interactions using non-brain specific endothelial and stromal cells promote gene regulation patterns that favor migration over proliferation.^[13a]^ Globally decreased KI67 expression also suggests an increased presence of quiescent GBM cells in perivascular niche models containing perivascular stromal cells. Quiescent tumor cells have been implicated in tumor re-growth and recurrence post-therapy,^[45]^ and the perivascular microenvironment has been shown to promote tumor cell dormancy in other cancer types.^[9a]^ Recent studies demonstrate that quiescent glioblastoma cells show heightened invasion potential and upregulate genes related to extracellular matrix interactions,^[46]^ consistent with phenotypic signature observed here. While KI67 expression experiments revealed differences in the abundance of proliferative cells, results from an EdU pulse experiment were able to identify the rate of tumor cell cycling events as a function of perivascular microenvironment.

Strikingly, our results revealed that a higher percentage of actively proliferating (KI67+) tumor cells incorporated EdU in the presence of perivascular stromal cells. Thus, we demonstrate that perivascular niche environments containing pericytes and astrocytes support the co-existence of discrete populations of slow and fast-cycling GBM cells, a phenotype which may facilitate short-term tumor growth and long-term disease persistence. Furthermore, the co-existence of quiescent and cycling tumor cells within our model better reflects the cell cycle heterogeneity found in *in vivo* glioblastoma specimens compared to *in vitro* models containing only tumor cells.^[47]^ Equivalent studies are largely intractable *in vivo*, highlighting the potential for complex multicellular perivascular models containing patient-derived specimens for gaining improved insight about how the cellular and microstructural environment in the perivascular niche may shape the evolution of a complex glioblastoma cell phenotype.

As the perivascular niche has been associated with chemoresistance,^[13b, 25a]^ we investigated the contributions of endothelial and perivascular stromal cells on GBM tumor cell phenotype after temozolomide treatment. In the absence of chemotherapy, pericytes and astrocytes generate a perivascular niche environment that inherently protects tumor cells from apoptosis (decreased cPARP+ fraction). EdU incorporation was reduced in tumor cells upon exposure to temozolomide, and this reduction in cycling rate is consistent with the ability of temozolomide to initiate G2/M cell cycle arrest, although whether cell cycle arrest results in DNA damage repair and continued proliferation, senescence, or apoptosis is unclear.^[48]^ We observed that cPARP and KI67 expression increased in response to TMZ exposure regardless of the presence of vascular cells, suggesting that temozolomide initiated apoptosis in a subfraction of tumor cells while also increasing the number of tumor cells in a proliferative state. In other cancer types such as breast cancer, patients with increased KI67 expression post-treatment have been correlated to worse outcomes compared to those with decreased or low KI67 expression.^[49]^ We additionally calculated apoptosis-to-proliferation ratios to compare chemosensitivity^[23, 50]^ as a result of perivascular co-culture and observed a significant increase in cPARP+/KI67+ cells in response to temozolomide treatment, but only in tumor-only samples. This data suggests that signaling from perivascular models provides a chemoprotective environment to glioblastoma tumor cells, though an explicit benefit of endothelial cells versus perivascular stromal (pericyte, astrocyte) was not observed. These studies are largely consistent with our prior findings using glioblastoma cell models and first-generation perivascular models comprised of non-brain specific endothelial and stromal cells, which showed the presence of perivascular signals, either from direct co-culture^[13a]^ or conditioned media,^[13b]^ may have a protective effect from TMZ treatment. While expanded cell cycle analysis and efforts to define the fraction of senescent, apoptotic, and DNA damage repair markers may ultimately provide an avenue to more fully characterize the effect of the perivascular niche in modulating glioblastoma response to therapy, hydrogel perivascular models containing perivascular stromal cells offers a framework to build a precision-medicine pipeline to consider a broader range of therapeutic interventions including emerging chemotherapy, radiotherapy, and antibody-drug-conjugate strategies.

## 4. Conclusion

In conclusion, we established a three-dimensional hydrogel model of the brain perivascular environment via self-assembly of human brain microvascular networks in GelMA hydrogels in order to investigate the role of the perivascular niche in glioblastoma progression.

Spontaneously forming microvascular networks demonstrate brain-relevant architectures, while the inclusion of perivascular stromal cells significantly increasing gene expression and functional markers related to tight junctions, basement membrane deposition, and membrane transporters. Notably, pericytes and astrocytes are critical for establishing a perivascular niche environment that enhances GBM cell invasion and limits tumor cell apoptosis. Perivascular networks containing pericytes and astrocytes enable the co-existence of quiescent and rapidly-cycling tumor cell populations with different spatial distributions. Finally, the engineered perivascular niche environment potentially mitigates tumor cell sensitivity to temozolomide. Overall, this platform provides a benchtop model to dissect the contributions of perivascular-derived cellular, biochemical, and biophysical cues on glioblastoma tumor progression, as well as a pre-clinical model for assessing patient-specific phenotypes and screening potential therapeutics in the context of the perivascular microenvironment.

## 5. Experimental Section/Methods

### Cell Culture

GBM6 and GBM12 cells were provided by Dr. Jann Sarkaria (Mayo Clinic, Rochester, MN) and cultured using the neurosphere method. GBM6 and GBM12 are derived from male patients, have a classical subtype, and demonstrate *EGFR* amplification. *MGMT* is unmethylated in GBM6 and methylated in GBM12. Briefly, cells were cultured in non-adherent culture flasks (50,000 cells/mL) in Knockout DMEM/F12 (Thermo Fisher Scientific, Waltham, MA) supplemented with StemPro Neural Supplement (Thermo Fisher Scientific, Waltham, MA), L-Glutamine (4 mM, Corning, Tewksbury, MA), EGF (20 ng/mL, Peprotech, Rocky Hill, NJ), bFGF (20 ng/mL, Peprotech, Rocky Hill, NJ), penicillin/streptomycin, and plasmocin prophylactic (Invivogen, San Diego, CA). Collectively, this media formulation will be referred to as neural stem cell (NSC) media. NSC media was replaced every two days.

Neurospheres were passaged when they were 100 - 200 µm in diameter (three days of culture for GBM12, seven days for GBM6). Neurospheres were dissociated using mechanical pipetting and TrypLE Express (Thermo Fisher Scientific, Waltham, MA) and re-plated on laminin-coated (Sigma Aldrich, St. Louis, MO) flasks for lentiviral transduction. MISSION® pLKO.1-puro-CMV-TagRFP™ positive control transduction particles (Sigma Aldrich, St. Louis, MO) were used to transduce GBM cells with RFP using a multiplicity of infection of 5. GBM cells were cultured with lentiviral particles for 18 hours, and then media was replaced and cells were cultured for an additional 48 hours before use. Transduction efficiency was approximately 90% as measured using fluorescence activated cell sorting (data not shown).

Differentiated GBM cells were obtained by plating cells derived from neurosphere culture on adherent flasks and culturing for two weeks in DMEM + 10% FBS.

Human brain microvascular endothelial cells (EC, Cell Systems, Kirkland, WA) were cultured on Attachment Factor-coated (Cell Systems) flasks in Endothelial Growth Medium 2 (EGM2, Lonza, Walkersville, MD) supplemented with plasmocin prophylactic. Human brain vascular pericytes (PC) and normal human astrocytes (AC), both from Sciencell (Carlsbad, CA), were cultured on poly-L-lysine (Sigma Aldrich, St. Louis, MO) coated flasks in Pericyte Growth Medium and Astrocyte Growth Medium (Sciencell) respectively. Astrocytes, pericytes, and endothelial cells were used before passages 3, 3, and 5 respectively. All cells were cultured at 37 °C and 5% CO2.

### Methacrylamide-Functionalized Gelatin (GelMA) Synthesis

GelMA was synthesized as previously described.^[31a]^ Briefly, porcine gelatin type A, 300 bloom (Sigma Aldrich, St. Louis, MO) was dissolved in PBS at 60 °C. 125 µL methacrylic anhydride (Sigma Aldrich, St. Louis, MO) was added dropwise per gram of gelatin, and the reaction proceeded for 1 hour with vigorous stirring (400 RPM). The reaction was quenched with PBS and dialyzed for seven days against deionized water with daily exchange. The product was then frozen and lyophilized. ^1^HNMR was used to determine the degree of functionalization (DOF). GelMA with DOF between 55 and 65% (data not shown) was used in this study.

### GBM Spheroid Formation

To form GBM6 spheroids for invasion studies, transduced GBM6 cells were dissociated using TrypLE Express and added to 96-well spheroid low-attachment plates (5000 cells/well) (Corning). Plates were centrifuged and then incubated on a rotating platform (60 RPM) at 37 °C for 48 hours.

### Vascular Culture in GelMA Hydrogels

5 wt% GelMA was dissolved in PBS at 65 °C. Lithium acylphosphinate (LAP, 0.1% w/v) was added as a photoinitiator. For cultures containing only ECs, 1 × 10^6^ EC/mL was re-suspended in the GelMA pre-polymer solution. For EC:PC co-cultures, a 3:1 EC:PC ratio was used with 1 × 10^6^ EC/mL. For EC:PC:AC co-cultures, 3:1:1 EC:PC:AC was used with 1 × 10^6^ EC/mL. Pre-polymer solutions with suspended cells were pipetted into circular Teflon molds (5 mm diameter, 1 mm thick) and polymerized under UV light (λ = 365 nm, 5.69 mW/cm^2^, 30 s). Hydrogels were cultured in EGM2 supplemented with 50 ng/mL VEGF (Peprotech) for up to seven days.

### GBM-Vascular Co-Culture in GelMA Hydrogels

For invasion experiments, a single GBM6 spheroid was encapsulated in each hydrogel alongside ECs, EC:PC, or EC:PC:AC in the ratios described above. Hydrogels containing only GBM6 spheroids were used as a negative control. For all other experiments, 2.5 × 10^5^/mL transduced GBM6 or GBM12 cells were encapsulated in single-cell suspension in pre-polymer solutions, either alone or with ECs, EC:PC, or EC:PC:AC in the ratios described above. Hydrogels were cultured for up to seven days in 500 µL/hydrogel of 1:1 EGM2 + 50 ng/mL VEGF : NSC media. When conditioned media collection was desired, media was switched to Endothelial Basal Medium 2 (EBM2) + 2% FBS on Day 6, and hydrogels were cultured for 24 hours before media collection. Conditioned media was sterile filtered and stored at - 80 °C.

### Immunofluorescent Staining

2D-plated cells and hydrogels were fixed using formalin, permeabilized using 0.5% Tween 20, and blocked using 2% abdil or 5% donkey serum (Sigma Aldrich) containing 0.1% Tween 20 before staining with primary antibodies. Primary antibodies included neural stem cell markers such as NESTIN (1:100, Abcam, Cambridge, UK), SOX2 (1:1000, Abcam or 1:100 Thermo Fisher Scientific), OLIG2 (1:40, R&D Systems, Minneapolis, MN); CD31 (1:100 R&D Systems or 1:200 Agilent, Santa Clara, CA); tight junction markers ZO-1 (1:50, Thermo Fisher Scientific) and OCCLUDIN (1:100, Thermo Fisher Scientific); RFP (1:500, Thermo Fisher Scientific); laminin (1;200, Abcam); PDGFRβ (1:40, R&D Systems); cleaved-PARP (1:400, Cell Signaling Technology, Danvers, MA) and differentiated neural and glial markers GFAP (1:1000, Abcam), βIII-Tubulin (1:62.5, Abcam), and O1 (1:100, R&D Systems); and KI67 (1:500, Thermo Fisher Scientific). Alexa Fluor 488, 555, and 633 were used as secondary antibodies (1:500, Thermo Fisher Scientific). All antibodies were applied overnight at 4 °C, and cells and hydrogels were washed with PBS + 0.1% Tween 20 between antibody incubations. Hoescht (1:2000) was used as a nuclear marker.

### EdU Pulse and Staining

The Click-iT EdU Cell Proliferation Kit for Imaging (Thermo Fisher Scientific, Waltham, MA) was used to assess the fraction of EdU+ cells in hydrogel cultures.^[5b]^ On Day 6, half of the media (250 µL) was removed from each hydrogel culture and replaced with fresh 1:1 EGM2 + 50 ng/mL VEGF : NSC media containing 20 µM EdU, such that the final concentration of EdU was 10 µM per well. Hydrogels were incubated with EdU for 24 hours, after which the media was removed and hydrogels were fixed in formalin for 15 minutes. The click reaction to attach Alexa Fluor 488 to the alkyne-containing EdU was performed per manufacturer’s instructions between primary and secondary antibody staining steps described in the previous section.

### GBM Spheroid Invasion Imaging and Analysis

Invasion experiments were imaged daily to track outgrowth migration of RFP+ GBM6 cells from spheroids.^[35]^ A DMi8 Yokogawa W1 spinning disk confocal microscope with a Hamamatsu EM-CCD digital camera (Leica Microsystems, Buffalo Grove, IL) was used to acquire fluorescent images at 5x or 10x magnification. ImageJ (NIH, Bethesda, MD) was used to obtain radial distributions of normalized fluorescent intensity to profile invasion for each sample. Normalized fluorescent intensity is calculated by summing the intensity of each pixel around the perimeter of a circle with radius (r) and dividing the summation by the number of pixels at radius (r). Images were binarized by manual thresholding, and intensity was normalized to lie between 0 and 1. The “Radial Profile” plugin was used to plot fluorescent intensity as a function of radius (r), with r = 0 established as the centroid of the spheroid. The centroid of the spheroid was obtained using “Analyze Particles” on a binary image of the spheroid. Radial distributions were characterized by defining r25, r50, and r75 metrics, which are the radii at which 25%, 50%, and 75% of the maximum fluorescent intensity are observed respectively. Total fluorescent intensity was calculated by integrating the radial distribution over radii:

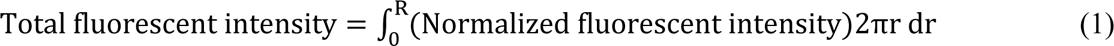

### Vessel Imaging and Metric Analysis

Using 10x magnification, z-stacks with an overall thickness of 200 µm and step size of 5 µm were obtained using the DMi8 to extract vessel metrics. Skeletons of the vascular networks were obtained using the TubeAnalyst macro (IRB Barcelona) in ImageJ. Skeletons were analyzed for metrics including total vessel length, average branch length, number of branches, and number of junctions using a Matlab code described by Crosby et al.^[51]^ Total vessel length, number of branches, and number of junctions were normalized to unit volume. Z-stacks with a step size of 2 µm were additionally obtained using 20x magnification to visualize tight junctions and interactions between astrocytes and endothelial cells. Z-stacks at 10x and 20x magnification were additionally obtained to visualize GBM - vascular co-cultures.

### SOX2/KI67/EdU/cPARP Imaging and Quantification

Using 10x magnification, z-stacks with an overall thickness of 200 µm and step size of 5 µm were obtained using the DMi8. Images were pre-processed in ImageJ by applying a background subtraction with a rolling ball radius of 10 – 15 µm as well as a contrast enhancement of 0.35. Processed images were analyzed using CellProfiler (Broad Institute, Boston, Massachusetts) to obtain the fractions of RFP labelled GBM cells that were positive for SOX2, KI67, EdU, or cPARP. Objects identified from SOX2, KI67, EdU, or cPARP staining were required to share at least 30% overlap in area with objects identified as GBM cells (using RFP intensity) in order to identify SOX2+, KI67+, EdU+, or cPARP+ GBM cells.

### Proximity Measurements between Tumor Cells and Vasculature

Maximum intensity projection images were used to analyze the proximity of tumor cells to the microvascular networks.^[11a, 31b, 52]^ CellProfiler was used to obtain separate binary images of SOX2- vs. SOX2+ tumor cells, KI67- vs. KI67+ tumor cells, and EdU- vs. EdU+ tumor cells from EC, EC:PC, and EC:PC:AC cultures. Intensity was adjusted to 0 for background and 1 for cells. Using ImageJ, images of microvascular networks were binarized via manual thresholding and a distance transform was performed. The distance transform produced images in which each pixel of the background was assigned a value equal to the distance of the pixel to the near microvascular structure. Images that underwent distance transformation were multiplied with the binarized images of tumor cells to obtain the distance of each tumor cell from the microvascular networks. Tumor cells were binned based on distance (0 – 10 μm, 10 – 20 μm, 20 – 30 μm, 30 – 40 μm, 40 – 50 μm, and >50 μm), and the fraction of tumor cells in each bin was calculated for each hydrogel. In line with prior literature,^[9a, 18]^ tumor cells within 50 μm of the microvascular network were defined as residing within the perivascular niche. Data was then averaged across all hydrogels in an experimental group.

### Temozolomide GR50 Determination

To determine the temozolomide (TMZ) dosage with which to perform experiments, we performed a dose-response experiment using GBM12-only (2.5×10^5^ cells/mL) hydrogels that had been cultured for seven days. Hydrogels were cultured with 0 – 600 µM temozolomide (TMZ, Calbiochem via Millipore Sigma, Burlington, MA), and 0.47% v/v DMSO (Sigma Aldrich) was used as a vehicle control. Hydrogels were cultured for 48 hours before viability was assessed using CellTiter-Glo 3D (Promega, Madison, WI). TMZ dosage and treatment period were informed by prior studies in our lab as well as chemotherapy studies performed in other three-dimensional *in vitro* models^[13, 53]^.

Hydrogels were equilibrated for 45 minutes at room temperature, and media was subsequently removed and replaced with 400 µL/hydrogel of 1:1 CellTiter-Glo Reagent : Knockout DMEM/F12 with StemPro Neural Supplement. Hydrogels were incubated at room temperature for one hour before the CellTiter:media solution was pipetted in triplicate into white-walled 96-well plates (Thermo Fisher Scientific) and luminescence was measured using a Biotek Synergy HT microplate reader (Winooski, VT). Viability was additionally assessed on untreated hydrogels at the beginning of the treatment period in order to calculate GR values according to Hafner et al. (**Figure S1**).^[22]^

### Temozolomide Treatment

Hydrogels containing GBM12 cells alone or GBM12 cells co-cultured with vascular cells were cultured for seven days as described in the previous sections. On Day 7, hydrogels were treated with 600 µM TMZ or a DMSO control (0.47% v/v) in 1 mL 1:1 EGM2 + 50 ng/mL VEGF : NSC media per hydrogel. Hydrogels were cultured for an additional 48 hours of treatment. On Day 9, hydrogels were fixed for KI67 and cPARP staining. An additional subset of hydrogels were maintained for another 24 hours for an EdU pulse. For these hydrogels, 750 µL of media was removed from each hydrogel culture and replaced with fresh 250 µL 1:1 EGM2 + 50 ng/mL VEGF : NSC media containing 20 µM EdU, such that the final concentration of EdU was 10 µM per well. Hydrogels were incubated with EdU for 24 hours, after which the media was removed and hydrogels were fixed for staining.

### RT-PCR

Vascular co-culture hydrogels were collected at Days 0, 3, and 7 and frozen at - 80 °C until further analysis.^[13a, 14]^ Hydrogels were crushed over dry ice using a pestle, and RNA was isolated using a RNeasy Plant Mini Kit (Qiagen, Hilden, Germany). RNA concentration was determined using a NanoDrop Lite spectrophotometer (Thermo Fisher Scientific, Waltham, MA). cDNA was obtained from reverse transcription by using a QuantiTect Reverse Transcription Kit (Qiagen, Hilden, Germany). RT-PCR reactions were prepared using Taqman Gene Expression Assays (**Table 1**) and Taqman Fast Advanced Master Mix (Thermo Fisher Scientific, Waltham, MA), and RT-PCR was performed using a QuantStudio 7 real-time PCR machine (Thermo Fisher Scientific, Waltham, MA). Data is expressed as a fold change calculated using the ΔΔCT method relative to Day 0 EC samples, with CD31 as an internal reference gene to normalize for the extent of vascularization between samples.

**Table 1.** Taqman Gene Expression Assays

| Gene | Assay ID |
| --- | --- |
| *PECAM1* | Hs01065279_m1 |
| *TJP1* | Hs01551861_m1 |
| *CLDN5* | Hs00533949_s1 |
| *OCLN* | Hs00170162_m1 |
| *LAMA4* | Hs00935293_m1 |
| *SLC2A1* | Hs00892681_m1 |

### Secretome Analysis

Conditioned media from GBM6 spheroid cultures with and without vascular cells was collected after seven days of culture as described previously.^[13b]^ The Proteome Profiler Human XL Cytokine Array (R&D Systems, Minneapolis, MN) was used to probe for differences in the secretome between experimental groups. Briefly, membranes were prepared according to manufacturer’s instructions and imaged using an ImageQuant LAS 4010 (Cytiva Life Sciences, Marlborough, MA) with an exposure time of two minutes.

Protein spot intensities were quantified using the MicroArray Profile plug-in (Optinav, Inc., Bellevue, WA) after background subtraction on ImageJ (NIH, Bethesda, MD). Intensities were normalized to the intensities of the positive reference spots in order to compare between membranes.

### Statistics

Statistics were performed using OriginPro (OriginLab, Northampton, MA). Normality of data was determined using the Shapiro-Wilk test, and equality of variance was determined using Levene’s test. For normal data, comparisons between paired and unpaired two groups were performed using a t-test, while comparisons between multiple groups were performed using a one-way ANOVA when assumptions were met. Tukey’s and Scheffe’s post-hoc tests were used for equal and unequal sample sizes respectively. In the case where data was not normal or groups had unequal variance, comparisons between two unpaired groups were performed using a Mann-Whitney test, comparisons between paired groups were performed using a Wilcoxon signed rank test, while comparisons between multiple groups were performed using a Kruskal-Wallis test with Dunn’s post-hoc. Significance was determined as p < 0.05. All quantitative analyses were performed on hydrogels set up across at least three independent experiments.

## Supporting Information

Supporting Information is available from the Wiley Online Library or from the author.

## Acknowledgements

The authors would like to thank Samantha Zambuto (UIUC), Alireza Sohrabi (UT-Austin), and Mark Schroeder (Mayo Clinic) for technical guidance that aided in experimental design and execution. The authors would also like to thank Zeng Hu (Mayo Clinic) for preparing and shipping primary tumor cells from patient-derived xenograft colonies. This work was supported by the National Cancer Institute of the National Institutes of Health (R01 CA256481, R01 CA197488) and the National Science Foundation Graduate Research Fellowship (DGE 1144245 to MTN). The content is solely the responsibility of the authors and does not necessarily represent the official views of the NIH. The authors also acknowledge additional funding provided by the Department of Chemical and Biomolecular Engineering and the Carl R. Woese Institute for Genomic Biology at the University of Illinois at Urbana-Champaign. The authors describe their contributions to the work according to the Contributor Roles Taxonomy (CRedit):^[54]^ M.T.N. – Conceptualization, Data curation, Formal Analysis, Visualization, Investigation, Methodology, Writing – original draft, Writing – review & editing. J.N.S – Resources, Writing – review & editing. B.A.C.H. - Conceptualization, Resources, Project administration, Funding acquisition, Supervision, Writing – review & editing.

## Supporting Information

**Figure S1.**
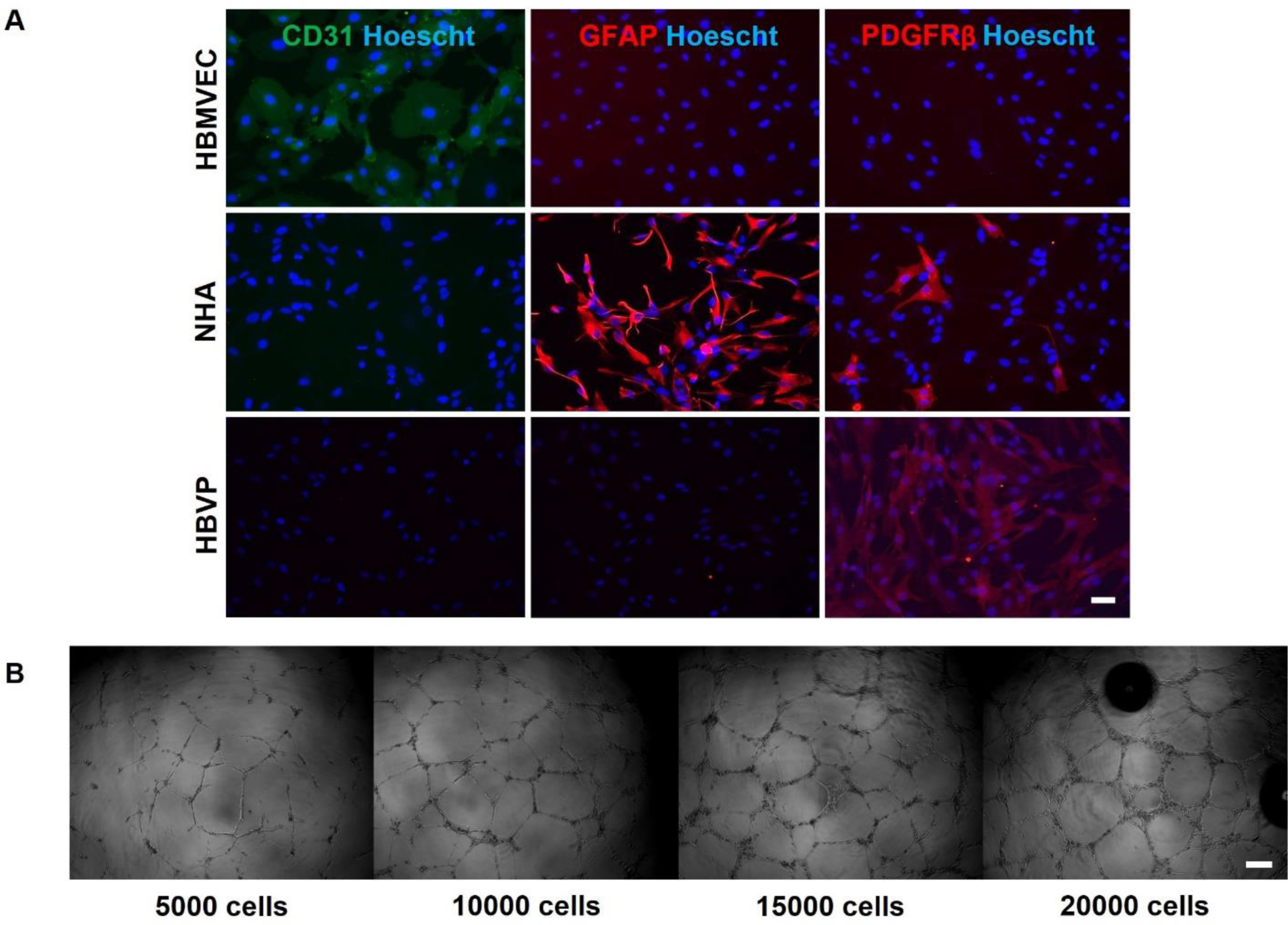
(A) Immunofluorescent markers for brain microvascular endothelial cells, astrocytes, and pericytes. Scale bar = 50 μm. **(B)** Matrigel tube formation assay for brain microvascular endothelial cells. Scale bar = 100 μm.

**Figure S2.**
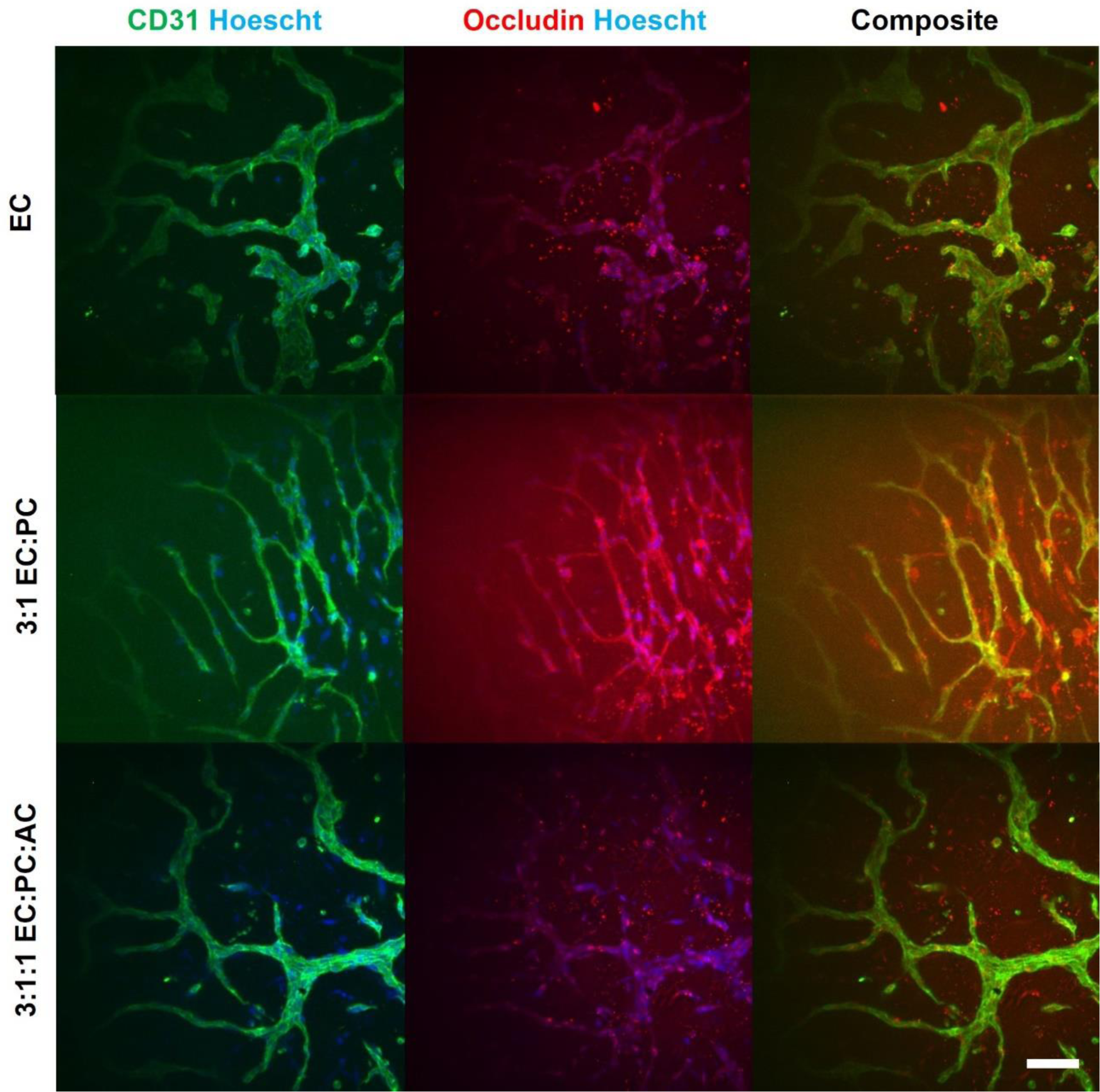
Immunofluorescent staining for occludins in microvascular cultures. Scale bar = 100 μm.

**Figure S3.**
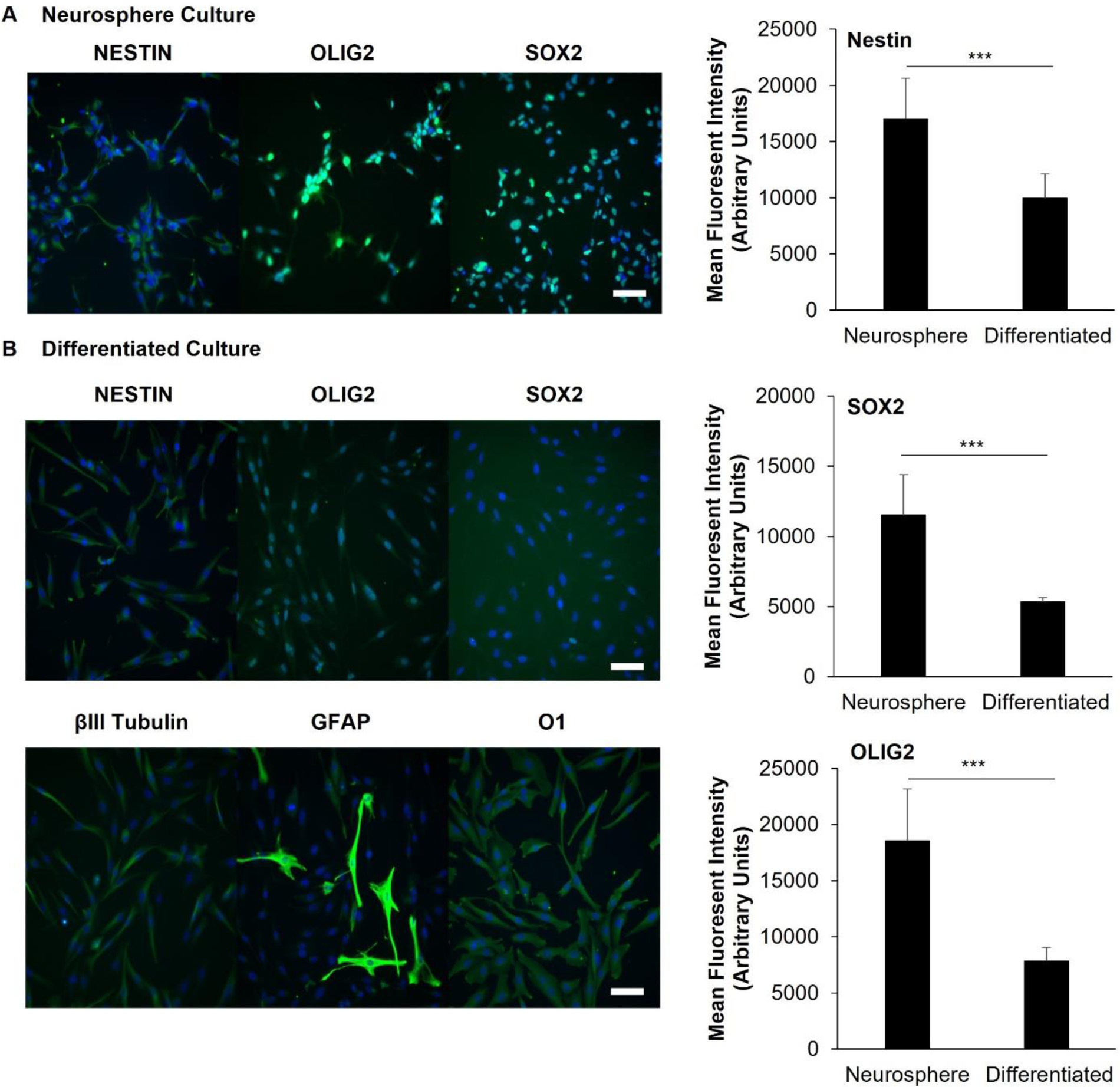
(A) Immunofluorescent staining for stem markers in GBM6 cells that have been grown in neurosphere culture and plated on laminin. Scale bar = 100 um. **(B)** Immunofluorescent staining for stem and differentiated markers for GBM6 cells grown in media containing FBS for two weeks. Scale bar = 100 μm. **(C)** Quantification of stem marker expression (assessed via mean fluorescent intensity) between neurosphere and differentiated cultures. ***p<0.001.

**Figure S4.**
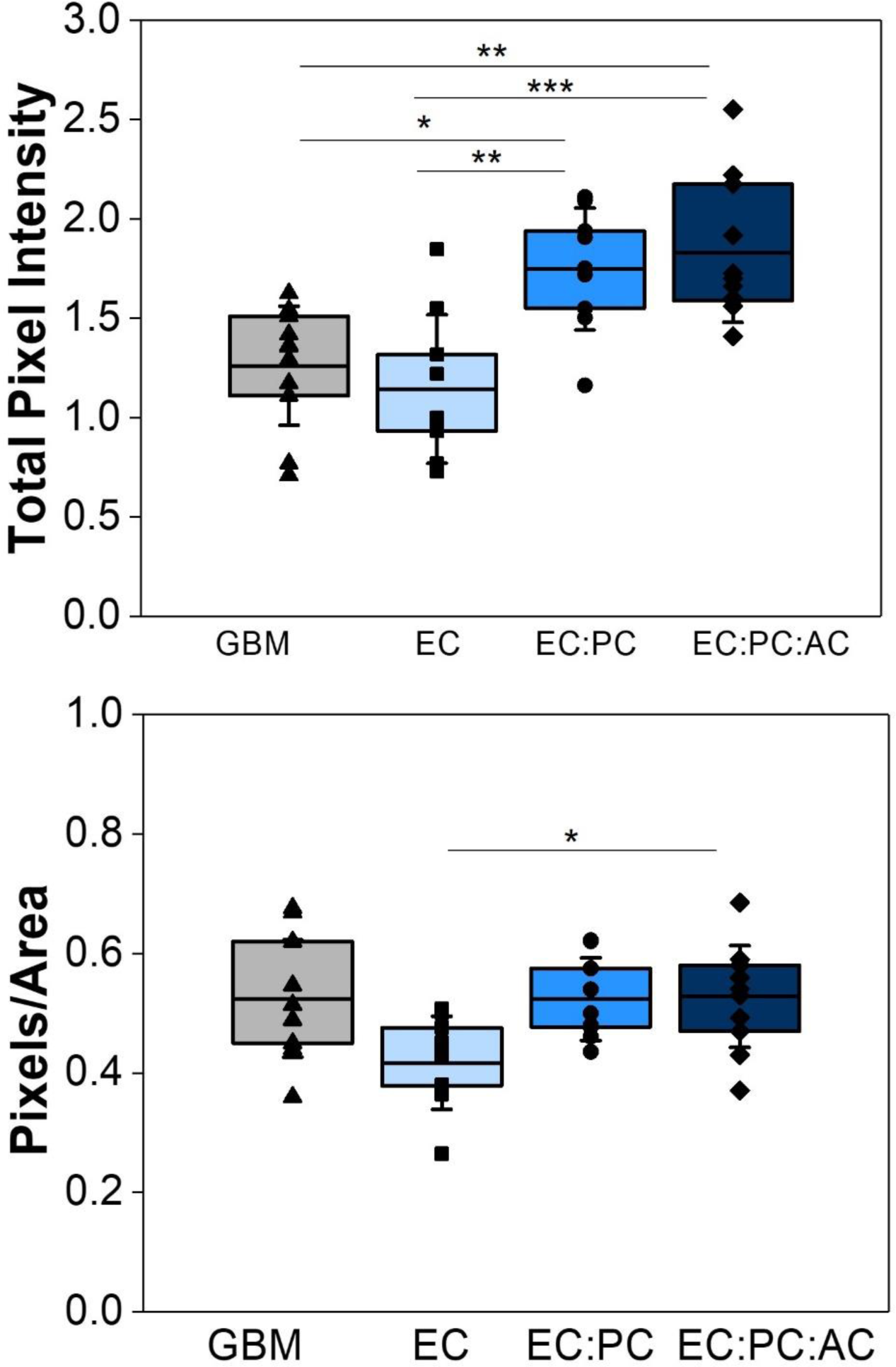
Quantification of total pixel intensity and total pixel intensity normalized to outgrowth area in spheroid invasion cultures. *p<0.05, **p<0.01, ***p<0.001, N = 9 – 11 hydrogels.

**Figure S5.**
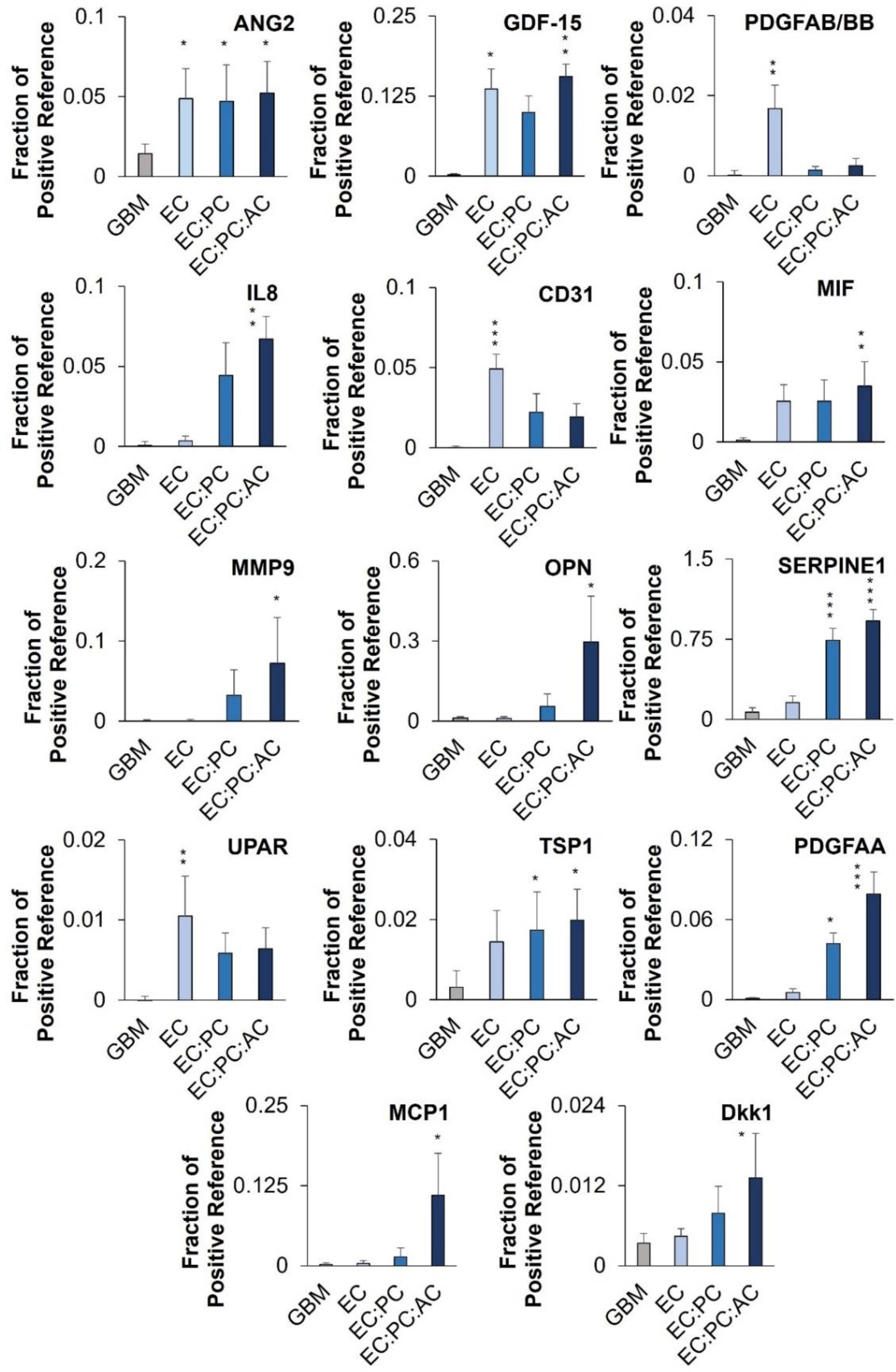
Quantitative comparison of proteins secreted by spheroid invasion cultures. *p<0.05, **p<0.01, ***p<0.001 compared to GBM6-only cultures; N = 5 conditioned media samples.

**Figure S6.**
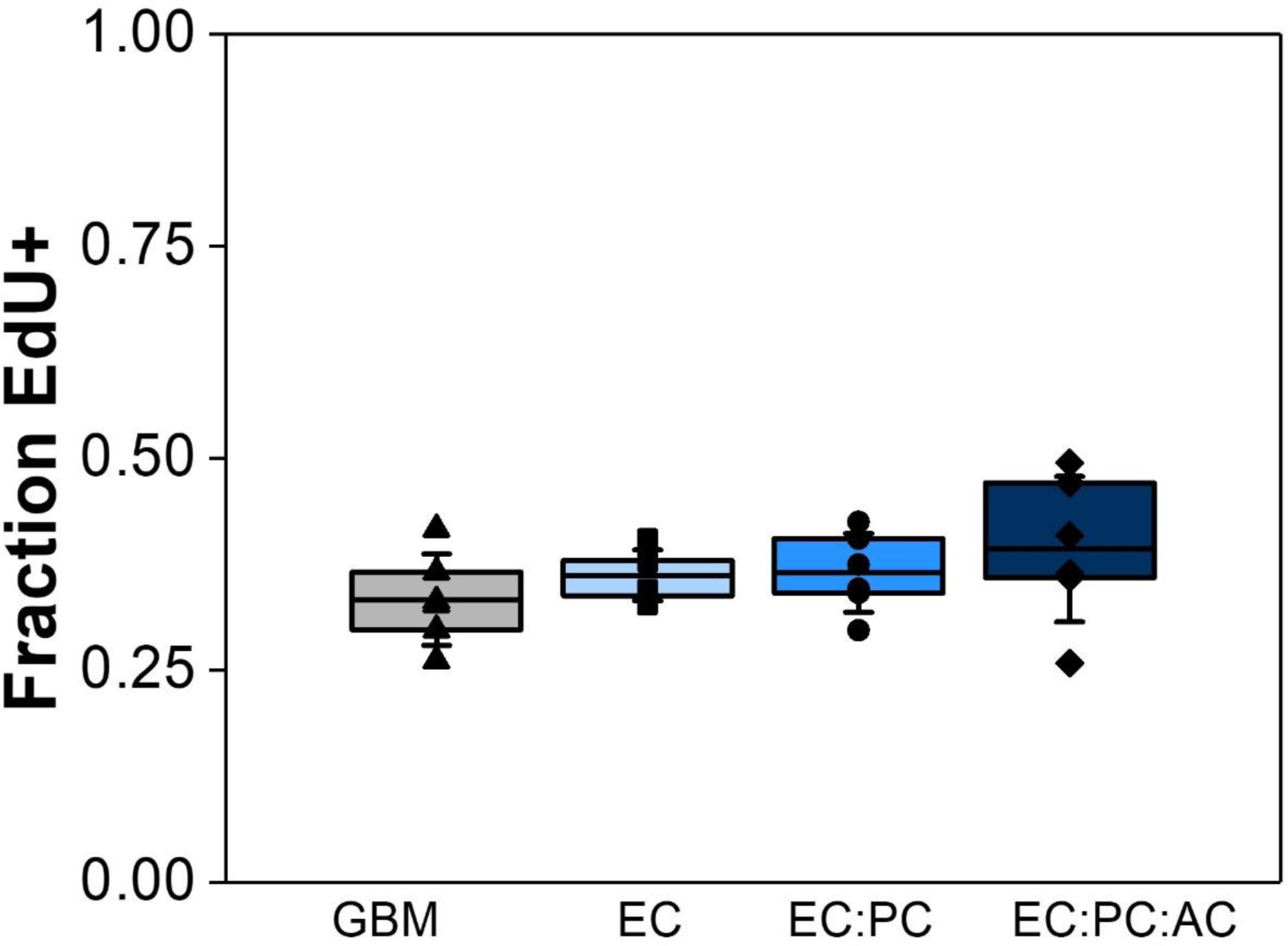
Quantification of the fraction of RFP-expressing GBM6 tumor cells that incorporate EdU in a 24-hour pulse. N = 6 hydrogels.

**Figure S7.**
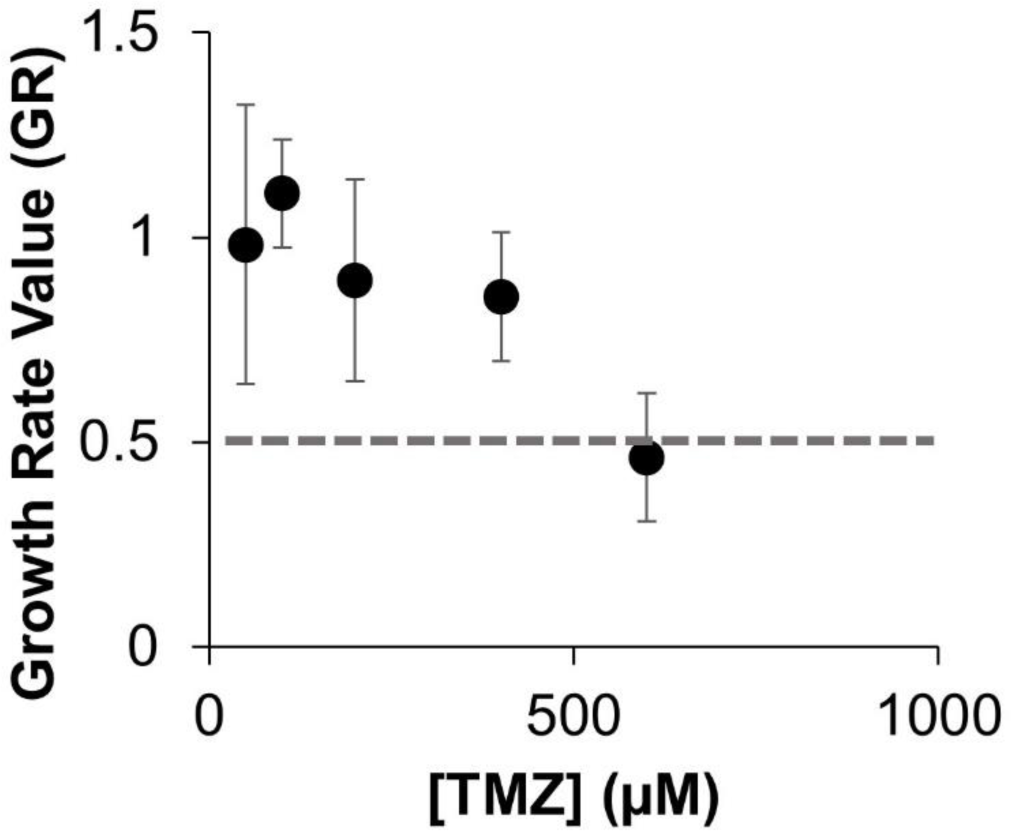
Dose-response curve for GBM12 cells treated with temozolomide for 48 hours. N = 4 - 6 hydrogels.

**Figure S8.**
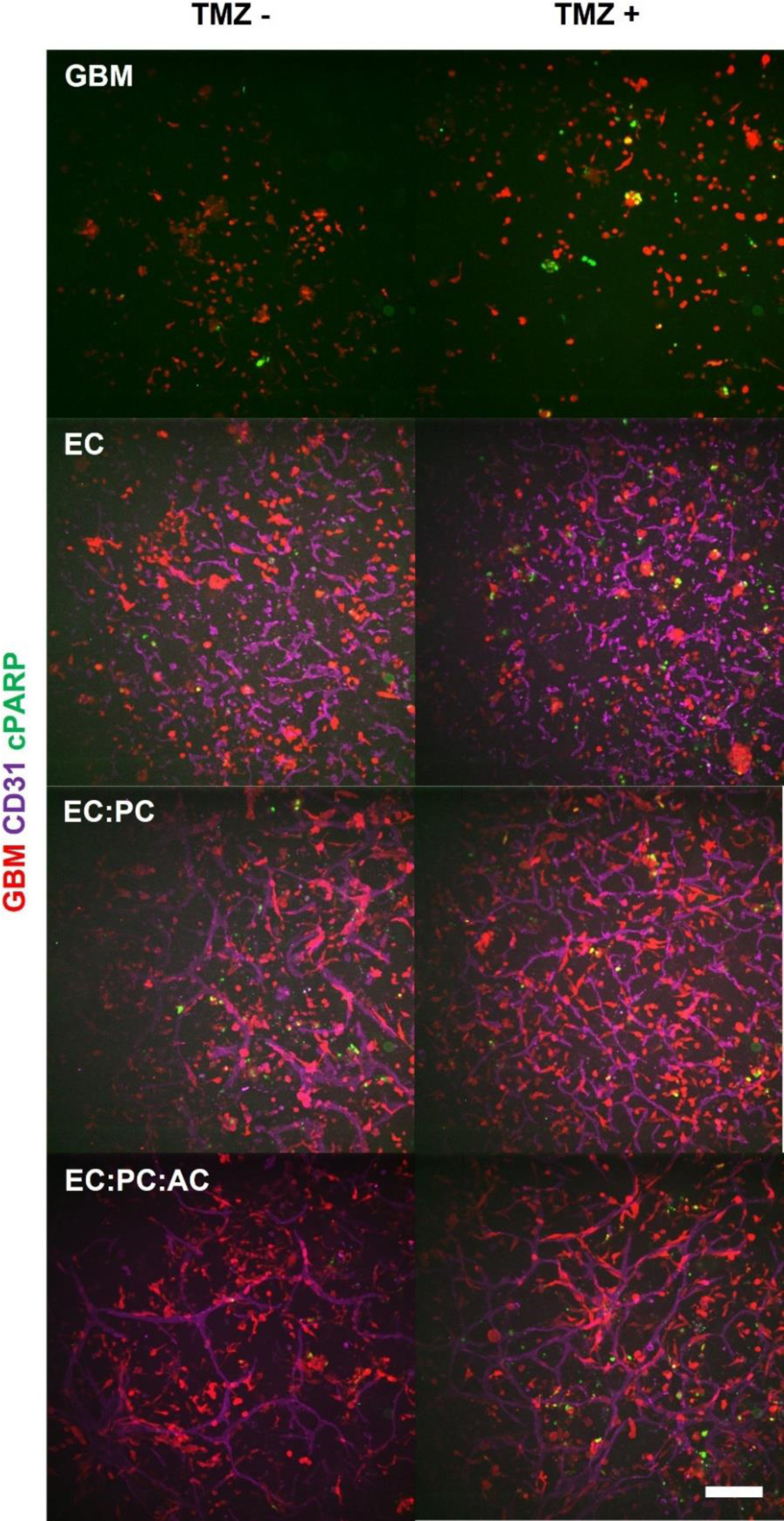
Representative images of tumor-vascular co-cultures with and without 48 hours of temozolomide treatment. cPARP staining was performed at the end of the treatment period. Scale bar = 200 μm.

**Figure S9.**
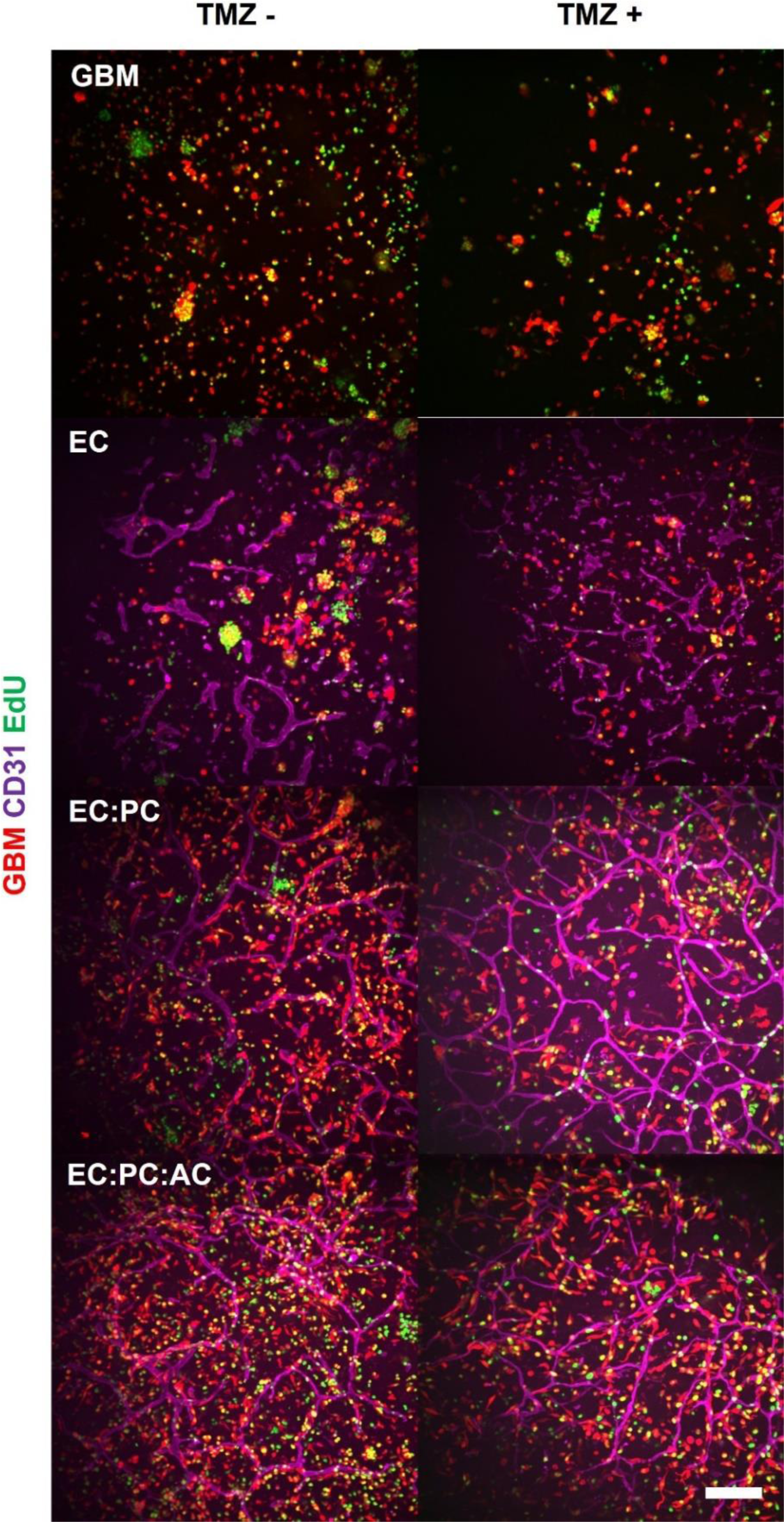
Representative images of tumor-vascular co-cultures with and without 48 hours of temozolomide treatment. EdU staining occurred after a 24-hour pulse of EdU after temozolomide treatment. Scale bar = 200 μm.

**Figure S10.**
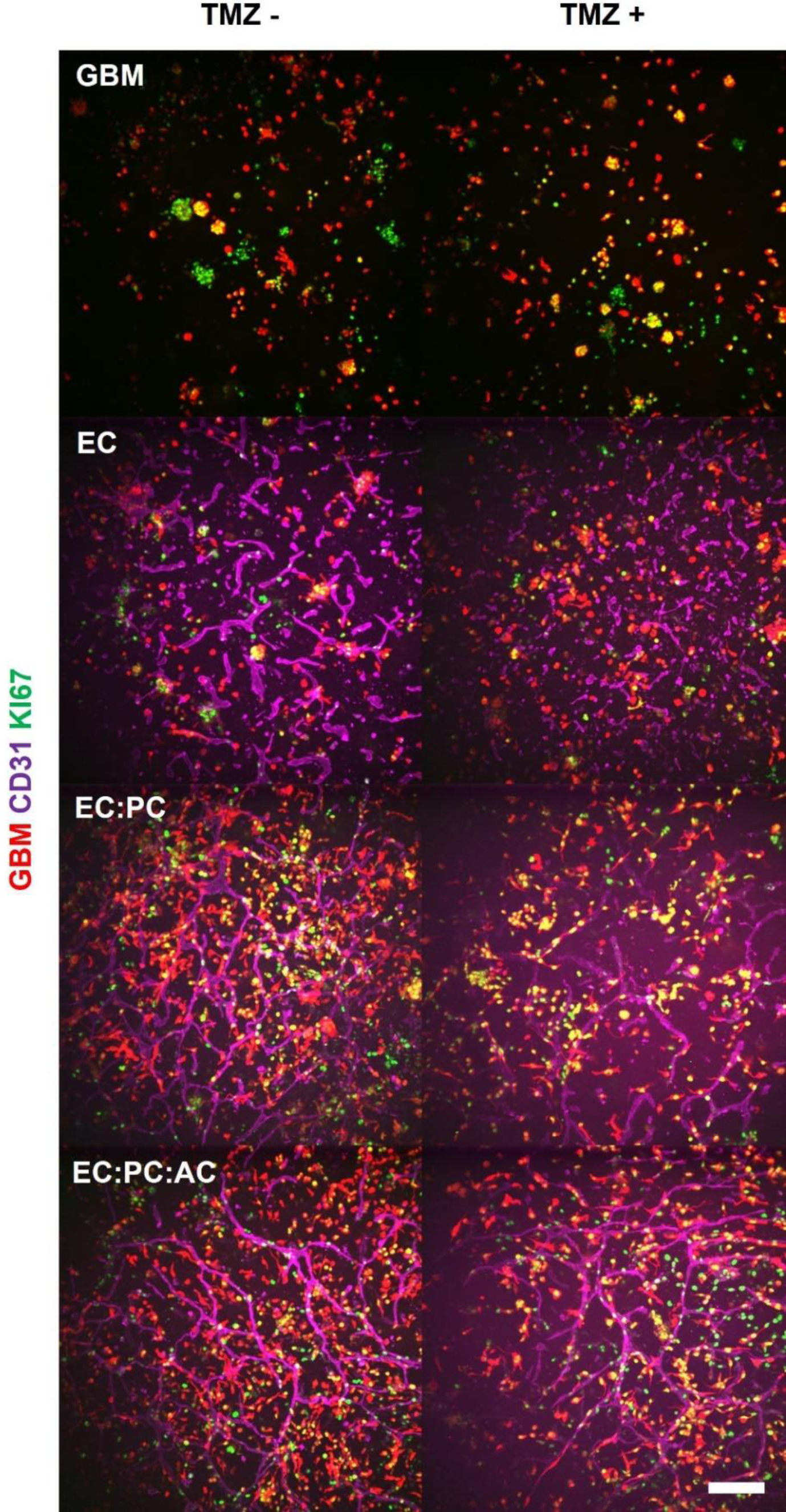
Representative images of tumor-vascular co-cultures with and without 48 hours of temozolomide treatment. KI67 staining was performed at the end of the treatment period. Scale bar = 200 μm.

## References

1. J. P. Thakkar, T. A. Dolecek, C. Horbinski, Q. T. Ostrom, D. D. Lightner, J. S. Barnholtz-Sloan, J. L. Villano, Cancer Epidemiol Biomarkers Prev 2014, 23, 1985.

2. L. M. DeAngelis, N Engl J Med 2001, 344, 114.

3. B. Campos, L. R. Olsen, T. Urup, H. S. Poulsen, Oncogene 2016, 35, 5819.

4. N. Cancer Genome Atlas Research, Nature 2008, 455, 1061 S. Darmanis, S. A. Sloan, D. Croote, M. Mignardi, S. Chernikova, P. Samghababi, Y. Zhang, N. Neff, M. Kowarsky, C. Caneda, G. Li, S. D. Chang, I. D. Connolly, Y. Li, B. A. Barres, M. H. Gephart, S. R. Quake, Cell Reports 2017, 21, 1399.

5. T. A. Ulrich, E. M. de Juan Pardo, S. Kumar, Cancer Research 2009, 69, 4167 W. Xiao, R. Zhang, A. Sohrabi, A. Ehsanipour, S. Sun, J. Liang, C. M. Walthers, L. Ta, D. A. Nathanson, S. K. Seidlits, Cancer Research 2018, 78, 1358 S. Pedron, G.L, Wolter, J.-W. E. Chen, S. E. Laken, J. N. Sarkaria, B. A. C. Harley, Biomaterials 2019, 219, 119371.

6. C. Calabrese, H. Poppleton, M. Kocak, T. L. Hogg, C. Fuller, B. Hamner, E. Y. Oh, L. W. Gaber, D. Finklestein, M. Allen, A. Frank, I. T. Bayazitov, S. S. Zakharenko, A. Gajjar, A. Davidoff, R. J. Gilbertson, Cancer Cell 2007, 11, 69.

7. A. Farin, S. O. Suzuki, M. Weiker, J. E. Goldman, J. N. Bruce, P. Canoll, Glia 2006, 53, 799 S. Watkins, S. Robel, I. F. Kimbrough, S. M. Robert, G. Ellis-Davies, H. Sontheimer, Nature communications 2014, 5, 4196.

8. J. D. Lathia, S. C. Mack, E. E. Mulkearns-Hubert, C. L. L. Valentim, J. N. Rich, Genes & Development 2015, 29, 1203.

9. C. M. Ghajar, H. Peinado, H. Mori, I. R. Matei, K. J. Evason, H. Brazier, D. Almeida, A. Koller, K. A. Hajjar, D. Y. Stainier, E. I. Chen, D. Lyden, M. J. Bissell, Nat Cell Biol 2013, 15, 807 Z. Cao, B.-S. Ding, P. Guo, Sharrell B. Lee, Jason M. Butler, Stephanie C. Casey, M. Simons, W. Tam, Dean W. Felsher, K. Shido, A. Rafii, Joseph M. Scandura, S. Rafii, Cancer Cell 2014, 25, 350.

10. G. J. Kitange, B. L. Carlson, M. A. Schroeder, P. T. Grogan, J. D. Lamont, P. A. Decker, W. Wu, C. D. James, J. N. Sarkaria, Neuro-Oncology 2009, 11, 281 S. K. Gupta, A. C. Mladek, B. L. Carlson, F. Boakye-Agyeman, K. K. Bakken, S. H. Kizilbash, M. A. Schroeder, J. Reid, J. N. Sarkaria, Clinical Cancer Research 2014, 20, 3730.

11. S. Rao, R. Sengupta, E. J. Choe, B. M. Woerner, E. Jackson, T. Sun, J. Leonard, D. Piwnica-Worms, J. B. Rubin, PLOS ONE 2012, 7, e33005 B. H. Rath, J. M. Fair, M. Jamal, K. Camphausen, P. J. Tofilon, PloS one 2013, 8, e54752 A. Mega, M. Hartmark Nilsen, L. W. Leiss, N. P. Tobin, H. Miletic, L. Sleire, C. Strell, S. Nelander, C. Krona, D. Hägerstrand, P. Ø. Enger, M. Nistér, A. Östman, Glia 2020, 68, 316.

12. L. J. Bray, M. Binner, A. Holzheu, J. Friedrichs, U. Freudenberg, D. W. Hutmacher, C. Werner, Biomaterials 2015, 53, 609 A. Sobrino, D. T. T. Phan, R. Datta, X. Wang, S. J. Hachey, M. Romero-López, E. Gratton, A. P. Lee, S. C. George, C. C. W. Hughes, Scientific Reports 2016, 6, 31589.

13. M. T. Ngo, B. A. C. Harley, Biomaterials 2019, 198, 122 M. T. Ngo, E. Karvelis, B. A. C. Harley, Integr Biol (Camb) 2020, 12, 139.

14. M. Campisi, Y. Shin, T. Osaki, C. Hajal, V. Chiono, R. D. Kamm, Biomaterials 2018, 180, 117.

15. J. N. Sarkaria, B. L. Carlson, M. A. Schroeder, P. Grogan, P. D. Brown, C. Giannini, K. V. Ballman, G. J. Kitange, A. Guha, A. Pandita, C. D. James, Clinical Cancer Research 2006, 12, 2264.

16. R. L. Bowman, Q. Wang, A. Carro, R. G. Verhaak, M. Squatrito, Neuro Oncol 2017, 19, 139.

17. R. M. R. Gangemi, F. Griffero, D. Marubbi, M. Perera, M. C. Capra, P. Malatesta, G. K. Ravetti, G. L. Zona, A. Daga, G. Corte, STEM CELLS 2009, 27, 40.

18. Y. Xiao, D. Kim, B. Dura, K. Zhang, R. Yan, H. Li, E. Han, J. Ip, P. Zou, J. Liu, A. T. Chen, A. O. Vortmeyer, J. Zhou, R. Fan, Advanced Science 2019, 6, 1801531.

19. A. Salic, T. J. Mitchison, Proceedings of the National Academy of Sciences 2008, 105, 2415.

20. K. Wang, F. M. Kievit, A. E. Erickson, J. R. Silber, R. G. Ellenbogen, M. Zhang, Adv Healthc Mater 2016, 5, 3173.

21. B. L. Carlson, J. L. Pokorny, M. A. Schroeder, J. N. Sarkaria, Current protocols in pharmacology / editorial board, S.J. Enna 2011, Chapter 14, Unit 14 16.

22. M. Hafner, M. Niepel, M. Chung, P. K. Sorger, Nat Methods 2016, 13, 521.

23. A. Dominijanni, M. Devarasetty, S. Soker, iScience 2020, 23, 101851 H. Kuriyama, K. R. Lamborn, J. R. O’Fallon, N. Iturria, T. Sebo, P. L. Schaefer, B. W. Scheithauer, J. C. Buckner, N. Kuriyama, R. B. Jenkins, M. A. Israel, Neuro-Oncology 2002, 4, 179.

24. R. R. Chhipa, Q. Fan, J. Anderson, R. Muraleedharan, Y. Huang, G. Ciraolo, X. Chen, R. Waclaw, L. M. Chow, Z. Khuchua, M. Kofron, M. T. Weirauch, A. Kendler, C. McPherson, N. Ratner, I. Nakano, N. Dasgupta, K. Komurov, B. Dasgupta, Nat Cell Biol 2018, 20, 823 A. Vartanian, S. K. Singh, S. Agnihotri, S. Jalali, K. Burrell, K. D. Aldape, G. Zadeh, Neuro Oncol 2014, 16, 1167 J.-W. Chen, S. Leary, V. Barnhouse, J. Sarkaria, B. A. C. Harley, Tissue Eng Part A 2021 J.-W. Chen, A. Blazek, J. Lumibao, H. R. Gaskins, B. A. C. Harley, Biomater Sci 2018, 6, 854.

25. T. Borovski, P. Beke, O. van Tellingen, H. M. Rodermond, J. J. Verhoeff, V. Lascano, J. B. Daalhuisen, J. P. Medema, M. R. Sprick, Oncogene 2013, 32, 1539 M. G. McCoy, D. Nyanyo, C. K. Hung, J. P. Goerger, W. R. Zipfel, R. M. Williams, M. Nishimura, C. Fischbach, Scientific Reports 2019, 9, 9069 D. Truong, R. Fiorelli, E. S. Barrientos, E. L. Melendez, N. Sanai, S. Mehta, M. Nikkhah, Biomaterials 2019, 198, 63.

26. A. Raza, M. J. Franklin, A. Z. Dudek, American Journal of Hematology 2010, 85, 593.

27. N. J. Abbott, L. Ronnback, E. Hansson, Nature reviews. Neuroscience 2006, 7, 41.

28. B. H. Rath, A. Wahba, K. Camphausen, P. J. Tofilon, Cancer Medicine 2015, 4, 1705.

29. M. E. Katt, E. V. Shusta, Current Opinion in Chemical Engineering 2020, 30, 42.

30. A. Al Ahmad, C. B. Taboada, M. Gassmann, O. O. Ogunshola, J Cereb Blood Flow Metab 2011, 31, 693.

31. J.-W. E. Chen, S. Pedron, B. A. C. Harley, Macromolecular Bioscience 2017, 17, 1700018 M. T. Ngo, B. A. Harley, Adv Healthc Mater 2017, 6.

32. K. Haase, G. S. Offeddu, M. R. Gillrie, R. D. Kamm, Advanced Functional Materials 2020, 30, 2002444.

33. A. Giese, R. Bjerkvig, M. E. Berens, M. Westphal, Journal of Clinical Oncology 2003, 21, 1624.

34. C. J. Liu, G. A. Shamsan, T. Akkin, D. J. Odde, Biophysical Journal 2019, 117, 1179.

35. J. E. Chen, S. Pedron, P. Shyu, Y. Hu, J. N. Sarkaria, B. A. C. Harley, Front Mater 2018, 5.

36. M. G. McCoy, D. Nyanyo, C. K. Hung, J. P. Goerger, R. Z. W. R. M. Williams, N. Nishimura, C. Fischbach, Sci Rep 2019, 9, 9069.

37. G. Bergers, S. Song, Neuro Oncol 2005, 7, 452.

38. R. M. Herrera-Perez, S. L. Voytik-Harbin, J. N. Sarkaria, K. E. Pollok, M. L. Fishel, J. L. Rickus, *PLoS One* 2018, 13, e0194183.

39. J. Holash, P. C. Maisonpierre, D. Compton, P. Boland, C. R. Alexander, D. Zagzag, G. D. Yancopoulos, S. J. Wiegand, Science 1999, 284, 1994 T. Kawataki, H. Naganuma, A. Sasaki, H. Yoshikawa, K. Tasaka, H. Nukui, Neuropathology 2000, 20, 161 S. Kono, J. S. Rao, J. M. Bruner, R. Sawaya, J Neuropathol Exp Neurol 1994, 53, 256.

40. P. Codó, M. Weller, K. Kaulich, D. Schraivogel, M. Silginer, G. Reifenberger, G. Meister, P. Roth, Oncotarget 2016, 7, 7732 D. W. Infanger, Y. Cho, B. S. Lopez, S. Mohanan, S. C. Liu, D. Gursel, J. A. Boockvar, C. Fischbach, Cancer Res 2013, 73, 7079 C. Lindemann, V. Marschall, A. Weigert, T. Klingebiel, S. Fulda, Neoplasia 2015, 17, 481 C. Piette, M. Deprez, T. Roger, A. Noël, J.-M. Foidart, C. Munaut, Journal of Biological Chemistry 2009, 284, 32483 D.-Y. Lu, W.-L. Yeh, S.-M. Huang, C.-H. Tang, H.-Y. Lin, S.-J. Chou, Neuro-Oncology 2012, 14, 1367 H. Feng, K. W. Liu, P. Guo, P. Zhang, T. Cheng, M. A. McNiven, G. R. Johnson, B. Hu, S. Y. Cheng, Oncogene 2012, 31, 2691 T. Daubon, C. Léon, K. Clarke, L. Andrique, L. Salabert, E. Darbo, R. Pineau, S. Guérit, M. Maitre, S. Dedieu, A. Jeanne, S. Bailly, J.-J. Feige, H. Miletic, M. Rossi, L. Bello, F. Falciani, R. Bjerkvig, A. Bikfalvi, Nature Communications 2019, 10, 1146.

41. S. A. Raithatha, H. Muzik, H. Muzik, N. B. Rewcastle, R. N. Johnston, D. R. Edwards, P. A. Forsyth, Neuro Oncol 2000, 2, 145 K. K. Veeravalli, C. Chetty, S. Ponnala, C. S. Gondi, S. S. Lakka, D. Fassett, J. D. Klopfenstein, D. H. Dinh, M. Gujrati, J. S. Rao, PLOS ONE 2010, 5, e11583 S. Kenig, M. B. Alonso, M. M. Mueller, T. T. Lah, Cancer Lett 2010, 289, 53.

42. M. T. Ngo, V. R. Barnhouse, A. E. Gilchrist, B. P. Mahadik, C. J. Hunter, J. N. Hensold, N. Petrikas, B. A. C. Harley, Advanced Functional Materials 2021, 31, 2101541.

43. T. Borovski, J. J. Verhoeff, R. ten Cate, K. Cameron, N. A. de Vries, O. van Tellingen, D. J. Richel, W. R. van Furth, J. P. Medema, M. R. Sprick, Int J Cancer 2009, 125, 1222 T. S. Zhu, M. A. Costello, C. E. Talsma, C. G. Flack, J. G. Crowley, L. L. Hamm, X. He, S. L. Hervey-Jumper, J. A. Heth, K. M. Muraszko, F. DiMeco, A. L. Vescovi, X. Fan, Cancer Res 2011, 71, 6061.

44. A. Giese, M. A. Loo, N. Tran, D. Haskett, S. W. Coons, M. E. Berens, Int J Cancer 1996, 67, 275.

45. J. Chen, Y. Li, T.-S. Yu, R. M. McKay, D. K. Burns, S. G. Kernie, L. F. Parada, Nature 2012, 488, 522 B. Campos, Z. Gal, A. Baader, T. Schneider, C. Sliwinski, K. Gassel, J. Bageritz, N. Grabe, A. von Deimling, P. Beckhove, C. Mogler, V. Goidts, A. Unterberg, V. Eckstein, C. Herold-Mende, The Journal of pathology 2014, 234, 23.

46. R. Tejero, Y. Huang, I. Katsyv, M. Kluge, J. Y. Lin, J. Tome-Garcia, N. Daviaud, Y. Wang, B. Zhang, N. M. Tsankova, C. C. Friedel, H. Zou, R. H. Friedel, EBioMedicine 2019, 42, 252 R. J. Atkins, S. S. Stylli, N. Kurganovs, S. Mangiola, C. J. Nowell, T. M. Ware, N. M. Corcoran, D. V. Brown, A. H. Kaye, A. Morokoff, R. B. Luwor, C. M. Hovens, T. Mantamadiotis, Experimental cell research 2019, 374, 353.

47. A. P. Patel, I. Tirosh, J. J. Trombetta, A. K. Shalek, S. M. Gillespie, H. Wakimoto, D. P. Cahill, B. V. Nahed, W. T. Curry, R. L. Martuza, D. N. Louis, O. Rozenblatt-Rosen, M. L. Suva, A. Regev, B. E. Bernstein, Science 2014, 344, 1396.

48. D. Aasland, L. Gotzinger, L. Hauck, N. Berte, J. Meyer, M. Effenberger, S. Schneider, E. E. Reuber, W. P. Roos, M. T. Tomicic, B. Kaina, M. Christmann, Cancer Res 2019, 79, 99 W. P. Roos, L. F. Z. Batista, S. C. Naumann, W. Wick, M. Weller, C. F. M. Menck, B. Kaina, Oncogene 2007, 26, 186 Y. Hirose, M. S. Berger, R. O. Pieper, Cancer Res 2001, 61, 1957.

49. G. von Minckwitz, W. D. Schmitt, S. Loibl, B. M. Müller, J. U. Blohmer, B. V. Sinn, H. Eidtmann, W. Eiermann, B. Gerber, H. Tesch, J. Hilfrich, J. Huober, T. Fehm, J. Barinoff, T. Rüdiger, E. Erbstoesser, P. A. Fasching, T. Karn, V. Müller, C. Jackisch, C. Denkert, Clinical Cancer Research 2013, 19, 4521 P. Cabrera- Galeana, W. Muñoz-Montaño, F. Lara-Medina, A. Alvarado-Miranda, V. Pérez-Sánchez, C. Villarreal-Garza, R. M. Quintero, F. Porras-Reyes, E. Bargallo-Rocha, I. Del Carmen, A. Mohar, O. Arrieta, Oncologist 2018, 23, 670.

50. T. W. Leung, W. C. Xue, A. N. Cheung, U. S. Khoo, H. Y. Ngan, Gynecologic oncology 2004, 92, 866.

51. C. O. Crosby, D. Valliappan, D. Shu, S. Kumar, C. Tu, W. Deng, S. H. Parekh, J. Zoldan, Tissue Engineering Part A 2019, 25, 746.

52. P. Carlson, A. Dasgupta, C. A. Grzelak, J. Kim, A. Barrett, I. M. Coleman, R. E. Shor, E. T. Goddard, J. Dai, E. M. Schweitzer, A. R. Lim, S. B. Crist, D. A. Cheresh, P. S. Nelson, K. C. Hansen, C. M. Ghajar, Nature Cell Biology 2019, 21, 238.

53. W. Xiao, S. Wang, R. Zhang, A. Sohrabi, Q. Yu, S. Liu, A. Ehsanipour, J. Liang, R D. Bierman, D. A. Nathanson, S. K. Seidlits, Matrix Biol 2020, 85–86, 128 E. A. Brooks, M. F. Gencoglu, D. C. Corbett, K. R. Stevens, S. R. Peyton, APL Bioeng 2019, 3, 026106.

54. L. Allen, J. Scott, A. Brand, M. Hlava, M. Altman, Nature 2014, 508, 312.

